# Putting the theory into ‘burstlet theory’: A biophysical model of bursts and burstlets in the respiratory preBötzinger complex

**DOI:** 10.1101/2021.11.19.469304

**Authors:** Ryan S. Phillips, Jonathan E. Rubin

## Abstract

Inspiratory breathing rhythms arise from synchronized neuronal activity in a bilaterally distributed brainstem structure known as the preBötzinger complex (preBötC). In *in vitro* slice preparations containing the preBötC, extracellular potassium must be elevated above physiological levels (to 7 − 9 *mM*) to observe regular rhythmic respiratory motor output in the hypoglossal nerve to which the preBötC projects. Reexamination of how extracellular *K*^+^ affects preBötC neuronal activity has revealed that low amplitude oscillations persist at physiological levels. These oscillatory events are sub-threshold from the standpoint of transmission to motor output and are dubbed burstlets. Burstlets arise from synchronized neural activity in a rhythmogenic neuronal subpopulation within the preBötC that in some instances may fail to recruit the larger network events, or bursts, required to generate motor output. The fraction of subthreshold preBötC oscillatory events (burstlet fraction) decreases sigmoidally with increasing extracellular potassium. These observations underlie the burstlet theory of respiratory rhythm generation. Experimental and computational studies have suggested that recruitment of the non-rhythmogenic component of the preBötC population requires intracellular *Ca*^2+^ dynamics and activation of a calcium-activated non-selective cationic current. In this computational study, we show how intracellular calcium dynamics driven by synaptically triggered *Ca*^2+^ influx as well as *Ca*^2+^ release/uptake by the endoplasmic reticulum in conjunction with a calcium-activated non-selective cationic current can explain all of the key observations underlying the burstlet theory of respiratory rhythm generation. Thus, we provide a mechanistic basis to unify the experimental findings on rhythm generation and motor output recruitment in the preBötC.

## Introduction

The complex neurological rhythms produced by central pattern generators (CPG) underlie numerous behaviors in healthy and pathological states. These activity patterns also serve as relatively experimentally accessible instances of the broader class of rhythmic processes associated with brain function. As such, CPGs have been extensively studied using a combination of experimental and computational approaches. The inspiratory CPG located in the preBötzinger complex (pre-BötC) in the mammalian respiratory brainstem is perhaps one of the most intensively investigated CPGs. Despite decades of research, the mechanisms of rhythm and pattern generation within this circuit remain unresolved and highly controversial; however, it appears that the pieces may now be in place to resolve this controversy.

Much of the debate in contemporary research into the mechanisms of preBötC rhythm and pat-tern generation revolves around the roles of specific ion currents, such as *I*_*NaP*_ and *I*_*CAN*_ (***Koizumi and Smith, 2008***; ***Thoby-Brisson and Ramirez, 2001***; ***Del Negro et al., 2002a***; ***Koizumi and Smith, 2008***; ***Koizumi et al., 2018***; ***Picardo et al., 2019***), and whether the observed rhythm is driven by an emergent network process (***Rekling and Feldman, 1998***; ***Del Negro et al., 2002b, 2005***; ***Rubin et al., 2009***; ***Sun et al., 2019***; ***Ashhad and Feldman, 2020***) and/or by intrinsically rhythmic or pacemaker neurons (***Johnson et al., 1994***; ***Koshiya and Smith, 1999***; ***Peña et al., 2004***). This debate is fueled by seemingly contradictory pharmacological blocking studies (***Del Negro et al., 2002a***; ***Peña et al., 2004***; ***Del Negro et al., 2005***; ***Pace et al., 2007b***; ***Koizumi and Smith, 2008***) and by new experimental studies (***Kam et al., 2013a***; ***Feldman and Kam, 2015***; ***Kallurkar et al., 2020***; ***Sun et al., 2019***; ***Ashhad and Feldman, 2020***) that challenge existing conceptual and computational models about the generation of activity patterns in the preBötC and underlie the so-called *burstlet theory* of respiratory rhythm generation.

The conceptual framework of burstlet theory posits that inspiratory oscillations arise from an emergent network process in a preBötC sub-population dedicated to rhythm generation and that a secondary pattern generating sub-population must be recruited in order to generate a full network burst capable of eliciting motor output. This hypothesis is supported by experimental preparations that compared local preBötC neuronal activity and motor output at the hypoglossal (XII) nerve in medullary slices. These studies found that in a low excitability state (controlled by the bath *K*^+^ concentration (*K*_*bath*_)), the preBötC generates a regular rhythm featuring a mixture of large and small amplitude network oscillations, dubbed *bursts* and *burstlets*, respectively, with only the bursts eliciting XII motor output (***Kam et al., 2013a***). Moreover, the fraction of low amplitude preBötC events (burstlet fraction) sigmoidally decreases with increasing *K*_*bath*_ and only a subset of preBötC neurons are active during burstlets (***Kallurkar et al., 2020***). Importantly, preBötC bursts can be blocked by application of cadmium (*Cd*^2+^), a calcium channel blocker, without affecting the ongoing burstlet rhythm (***Kam et al., 2013a***; ***Sun et al., 2019***), supporting the idea that rhythm generation occurs in a distinct preBötC subpopulation from pattern generation and demonstrating that conversion of a burst into a burstlet is a *Ca*^2+^-dependent process. Finally, rhythm generation in the burstlet population is hypothesized to result from an emergent network percolation process. This last idea was developed to explain holographic photostimulation experiments, which found that optically stimulating small subsets (4 − 9) of preBötC inspiratory neurons was sufficient to reliably evoke endogenous-like XII inspiratory bursts with delays averaging 255 ± 45 *ms* (***Kam et al., 2013b***). The small number of neurons required to evoke a network burst and the extended duration of the delays both differ from what would be predicted by existing computational preBötC models. Additionally, these delays are on a similar timescale to the ramping pre-inspiratory neuronal activity that precedes network-wide inspiratory bursts, leading to the hypothesis that stimulation of this small set of preBötC neurons kicks off an endogenous neuronal percolation process underlying rhythm generation, which could be initiated by the near-coincident spontaneous spiking of a small number of preBötC neurons.

The experimental underpinning of burstlet theory challenges current ideas about inspiratory rhythm and pattern generation. However, the proposed mechanisms of burst and burstlet generation remain hypothetical and, to date, there has not been a quantitative model that provides a unified, mechanistic explanation for the key experimental observations or that validates the conceptual basis for this theory. Interestingly, key components of burstlet theory, namely that inspiratory rhythm and pattern are separable processes and that large amplitude network-wide bursts depend on calcium-dependent mechanisms are supported by recent experimental and computational studies. Specifically, ***Koizumi et al. (2018***); ***Picardo et al. (2019***) showed that the amplitude (i.e. pattern) of preBötC and XII bursts is controlled, independently from the ongoing rhythm, by the transient receptor potential channel (TRPM4), a calcium-activated non-selective cation current (*I*_*CAN*_). These findings are consistent with burstlet theory, as they demonstrate that rhythm and pattern are separable processes at the level of the preBötC. Moreover, these experimental observations are robustly reproduced by a recent computational modeling study (***Phillips et al., 2019***), which shows that pattern generation can occur independently of rhythm generation. Consistent with burstlet theory, this model predicts that rhythm generation arises from a small subset of preBötC neurons, which in this model form a persistent sodium (*I*_*NaP*_) dependent rhythmogenic kernel, and that rhythmic synaptic drive from these neurons triggers post-synaptic calcium transients, *I*_*CAN*_ activation, and amplification of the inspiratory drive potential, which drives bursting in the rest of the network.

These recent results suggest that conversion of burstlets into bursts may be *Ca*^2+^ and *I*_*CAN*_ dependent, occurring when synaptically triggered calcium transients in non-rhythmogenic preBötC neurons are intermittently large enough for *I*_*CAN*_ activation to occur and to yield recruitment of these neurons into the network oscillation. The biophysical mechanism responsible for periodic amplification of *Ca*^2+^ transients is not known, however. In this computational study, we put to-gether and build upon these previous findings to show that periodic amplification of synaptically triggered *I*_*CAN*_ transients by calcium induced calcium release (CICR) from intracellular stores provides a plausible mechanism that can produce the observed conversion of burstlets into bursts and can explain all of the key observations underlying the burstlet theory of respiratory rhythm generation, thus providing a sound mechanistic basis for this conceptual framework.

## Results

### Calcium induced calcium release periodically amplifies intracellular calcium transients

Our first aim in this work was to test whether calcium induced calcium release from ER stores could repetitively amplify synaptically triggered *Ca*^2+^ transients. To address this aim, we constructed a cellular model that includes the endoplasmic reticulum. The model features a *Ca*^2+^ pump, which extrudes *Ca*^2+^ from the intracellular space, a sarcoendoplasmic reticulum calcium transport AT-Pase (SERCA) pump, which pumps *Ca*^2+^ from the intracellular space into the ER, and the *Ca*^2+^ activated inositol trisphosphate (IP3) receptor (Fig 1A). To simulate calcium transients synaptically generated from a rhythmogenic source (i.e., burstlets), we imposed a square wave *Ca*^2+^ current into the intracellular space with varied frequency and amplitude but fixed duration (250 *ms*) and we monitored the resulting intracellular *Ca*^2+^ transients. Depending on parameter values used, we observed various combinations of low and high amplitude *Ca*^2+^ responses, and we characterized how the fraction of *Ca*^2+^ transients that have low amplitude depends on values selected within the 2D parameter space parameterized by *Ca*^2+^ pulse frequency and amplitude. We found that the fraction of low amplitude *Ca*^2+^ transients decreases as either or both of the *Ca*^2+^ pulse frequency and amplitude are increased (Fig. 1B and example traces C1-C4).

**Figure 1.**
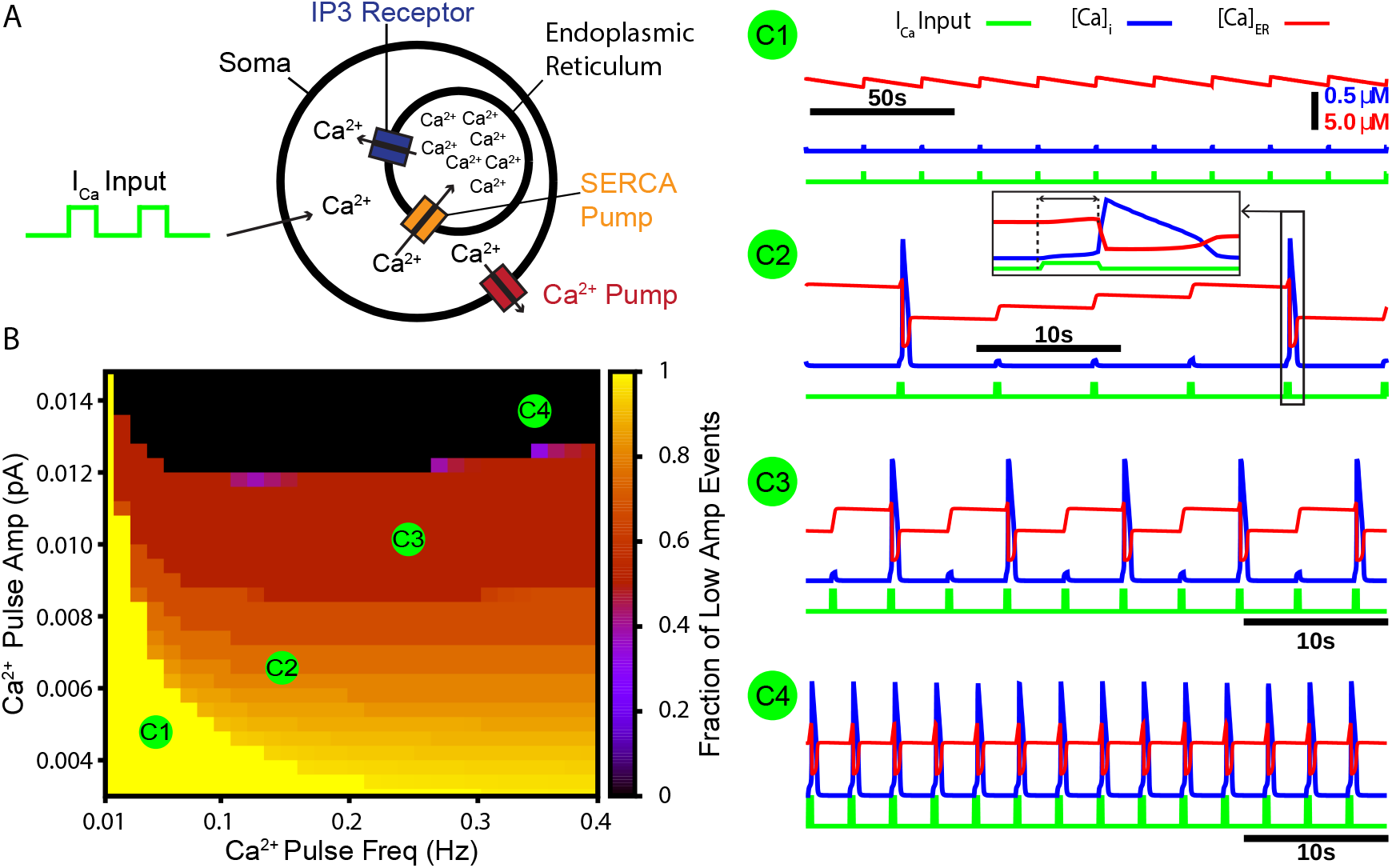
A periodic input in the form of a calcium current drives intermittent calcium induced calcium release from ER stores. (A) Schematic diagram of the model setup showing square wave profile of *Ca*^2+^ current input into the intracellular space, uptake of *Ca*^2+^ into the ER by the SERCA pump, *Ca*^2+^ release through the IP3 receptor, and extrusion of *Ca*^2+^ through a pump in the cell membrane. (B) Fraction of low amplitude intracellular *Ca*^2+^ transients as a function of the *Ca*^2+^ pulse frequency and amplitude. Pulse duration was fixed at 250 *ms*. (C1-C4) Example traces showing several ratios of low and high amplitude *Ca*^2+^ transients and the dynamics of the ER stores *Ca*^2+^ concentration. Inset in C2 highlights the delay between pulse onset and CICR. The pulse amplitude and frequency for each trace are indicated in panel B.

### Bursts and burstlets in a two-neuron preBötC network

Next we tested whether the CICR mechanism (Fig. 1) could drive intermittent recruitment in a reciprocally connected two neuron network that includes one intrinsically rhythmic and one non-rhythmic neuron, as a preliminary step towards considering the rhythm and pattern generating sub-populations of the preBötC suggested by burstlet theory (***Kam et al., 2013a***; ***Cui et al., 2016***; ***Kallurkar et al., 2020***; ***Ashhad and Feldman, 2020***) and recent computational investigation (***Phillips et al., 2019***). In this network, neuron 1 is an *I*_*NaP*_ -dependent intrinsically bursting neuron, with a burst frequency that is varied by injecting an applied current, *I*_*AP P*_ (Fig. 2 A2-A3). The rhythmic bursting from neuron 1 generates periodic postsynaptic currents (*I*_*Syn*_) in neuron 2, carried in part by *Ca*^2+^ ions, which are analogous to the square wave *Ca*^2+^ current in Fig 1. The amplitude of the postsynaptic *Ca*^2+^ transient is determined by the number of spikes per burst (Fig. 2A4) and by the parameter *P*_*SynCa*_, which determines the percentage of *I*_*Syn*_ carried by *Ca*^2+^ ions (see *materials and methods* for a full description of these model components). Conversion of a burstlet (isolated neuron 1 burst) into a network burst (recruitment of neuron 2) is dependent on CICR (see Fig. 2-Figure Supplement 1), which increases intracellular calcium above the threshold for *I*_*CAN*_ activation.

**Figure 2.**
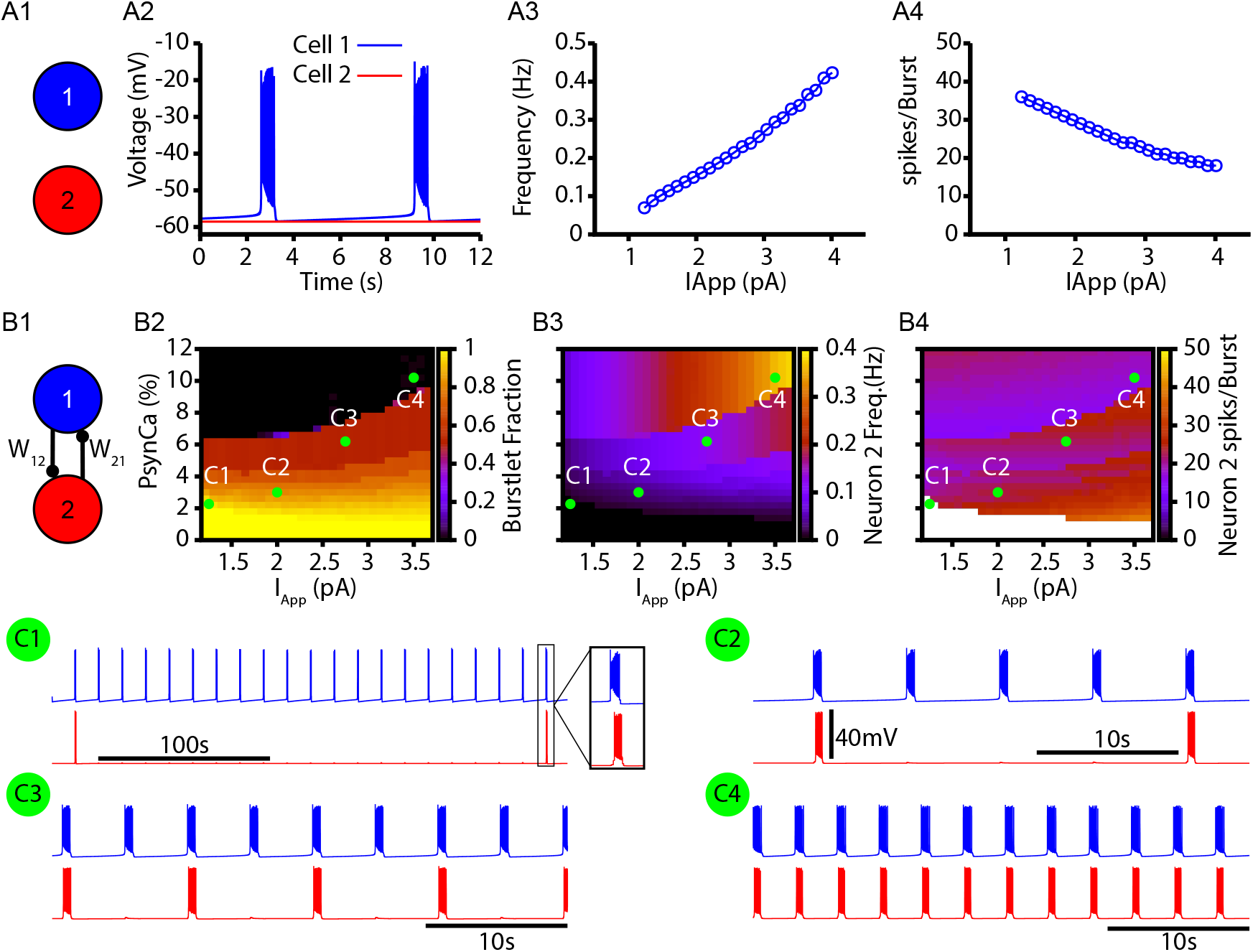
Bursts and burstlets in a two neuron preBötC network. (A1) Schematic diagram of the synaptically uncoupled network. The rhythm and pattern generating components of the network are represented by neuron 1 and 2, respectively. (A2) Example trace showing intrinsic bursting in neuron 1 and quiescence in neuron 2. (A3) Burst frequency and (A4) the number of spikes per burst in neuron 1 as a function of an applied current (*I*_*AP P*_). Neuron 2 remained quiescent within this range of *I*_*AP P*_. (B1) Schematic diagram of the synaptically coupled network. (B2-B4) 2D plots characterizing the (B2) burstlet fraction, (B3) neuron 2 (burst) frequency, and (B4) neuron 2 spikes per burst (burst amplitude) as a function of *I*_*AP P*_ and *P*_*SynCa*_. (C1-C4) Example traces for neuron 1 and 2 for various *I*_*AP P*_ and *P*_*SynCa*_ values indicated in (B2-B4). Notice the scale bar is 100 *s* in C1 and 10 *s* in C2-C4. Inset in C1 shows the burst shape not visible on the 100 *s* time scale. The model parameters used in these simulations are: (neuron 1 & 2) *K*_*Bath*_ = 8 *mM, g*_*Leak*_ = 3.35 *nS, W*_12_ = *W*_21_ = 0.006 *nS*; (Neuron 1) *g*_*NaP*_ = 3.33 *nS, g*_*CAN*_ = 0.0 *nS*, (Neuron 2) *g*_*NaP*_ = 1.5 *nS, g*_*CAN*_ = 1.5 *nS*.

In the reciprocally connected network, we first quantified the dependence of the burstlet fraction, which was defined as the number of burstlets (neuron 1 bursts without recruitment of neuron 2) divided by the total number of burstlets and network bursts (bursts in neuron 1 with recruitment of neuron 2), on *I*_*AP P*_ and *P*_*SynCa*_. Increasing *I*_*AP P*_ increases the burst frequency in neuron 1 and decreases the number of spikes per neuron 1 burst (Fig. 2A3,A4), consistent with past literature (***Butera et al., 1999***). These changes do not strongly impact the burstlet fraction until *I*_*AP P*_ grows enough, at which point the shorter, more rapid bursts of neuron 1 become less effective at recruiting neuron 2 and thus the burstlet fraction increases (Fig. 2B2). In general, increasing *P*_*SynCa*_ decreased the burstlet fraction (i.e., increased the frequency of neuron 2 recruitment) by causing a larger calcium influx with each neuron 1 burst; see Fig. 2B2 & C1-C4.

The burst frequency in neuron 2 is determined by the burst frequency of neuron 1 and the burstlet fraction. These effects determine the impact of changes in *P*_*SynCa*_ and *I*_*AP P*_ on neuron 2 burst frequency (Fig. 2 B3). As *I*_*AP P*_ increases, the rise in burstlet frequency implies that neuron 2 bursts in response to a smaller fraction of neuron 1 bursts, yet the rise in neuron 1 burst frequency means that these bursts occur faster. These two effects can balance to yield a relatively constant neuron 2 frequency, although the balance is imperfect and frequency does eventually increase. Increases in *P*_*SynCa*_ more straightforwardly lead to increases in neuron 2 burst frequency as the burstlet fraction drops.

Finally, the number of spikes per burst in neuron 2 is not strongly affected by changes in *I*_*AP P*_ and *P*_*SynCa*_ (Fig. 2 B4), suggesting an all-or-none nature of recruitment of bursting in neuron 2. Interestingly, the period between network bursts (i.e., time between neuron 2 recruitment events) can be on the order of hundreds of seconds (e.g., Fig. 2 C1). This delay is consistent with some of the longer timescales shown in experiments characterizing bursts and burstlets (***Kallurkar et al., 2020***).

### CICR supports bustlets and bursts in a data-constrained preBötC network model

Next, we tested whether the CICR mechanism presented in Figs.1 & 2 could underlie the conversion of burstlets into bursts in a larger preBötC model network including rhythm and pattern generating subpopulations and whether this network could capture the *K*_*bath*_-dependent changes in the burstlet fraction characterized in ***Kallurkar et al. (2020***). *K*_*bath*_ sets the extracellular *K*^+^ concentration, which in turn determines the driving force for any ionic currents that flux *K*^+^. In preBötC neurons these currents include the fast *K*^+^ current, which is involved in action potential generation, and the *K*^+^-dominated leak conductance, which primarily affects excitability (Fig. 3A). In our simulations, we modeled the potassium (_*K*_) and leak (_*Leak*_) reversal potentials as functions of *K*_*bath*_ using the Nernst and Goldman–Hodgkin–Katz equations. The resulting curves were tuned to match existing data from ***Koizumi and Smith (2008***), as shown in Fig. 3B. In our simulations, we found that intrinsic bursting is extremely sensitive to changes in *K*_*bath*_. However, with increasing *K*_*bath*_, intrinsic bursting could be maintained over a wide range of *K*^+^ concentrations when accompanied by increases in *g*_*Leak*_ (Fig. 3C). Additionally, the number of spikes per burst in the bursting regime increases with *K*_*bath*_ (Fig. 3 Figure Supplement 1). This *K*_*bath*_-dependence of *g*_*Leak*_ is consistent with experimental data showing that neuronal input resistance decreases with increasing *K*_*bath*_ (***Okada et al., 2005***).

**Figure 3.**
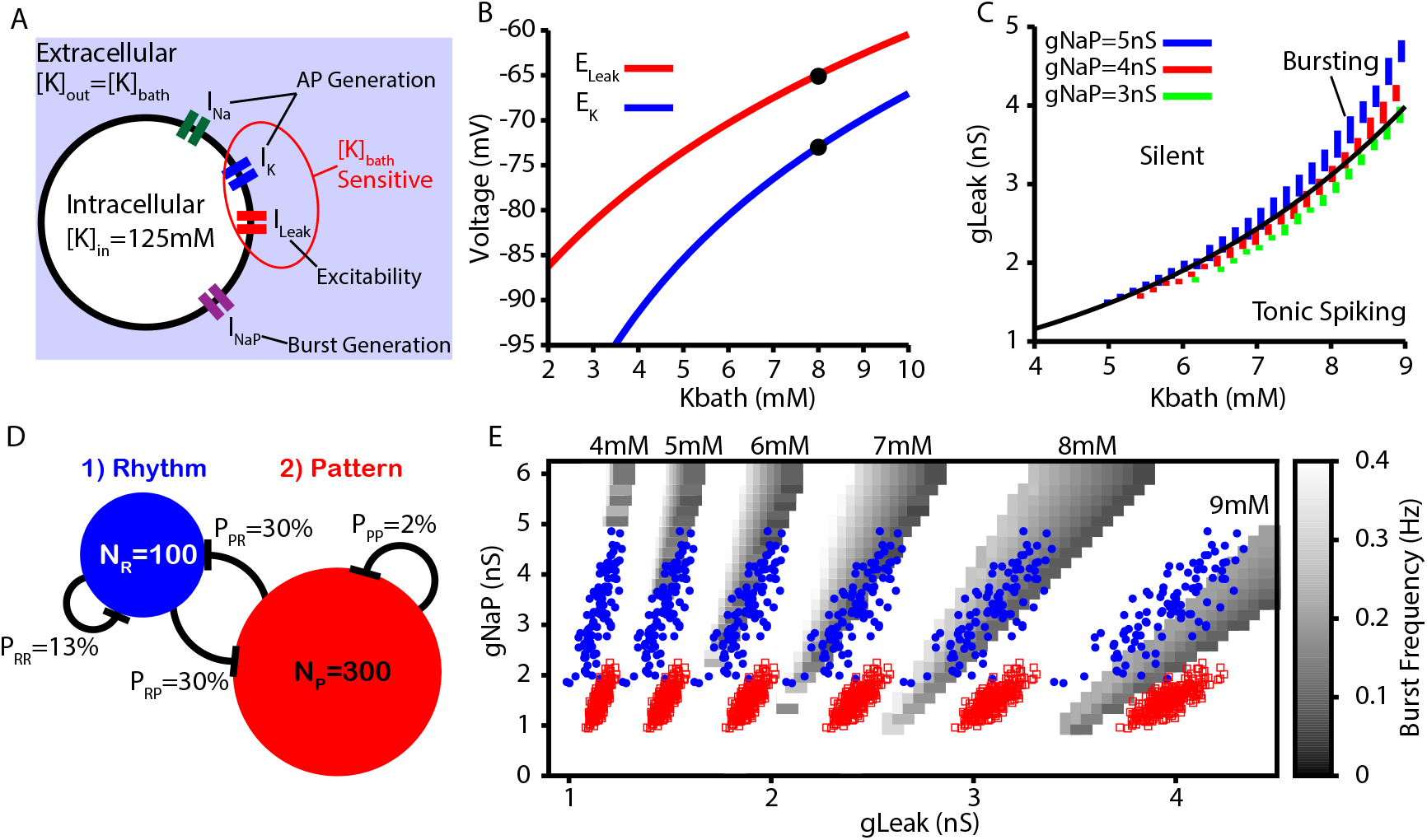
Intrinsic cellular and network dynamics depend on the bath potassium concentration. (A) Schematic diagram of an isolated model preBötC neuron showing the simulated ion channels involved in AP generation, excitability and burst generation as well as indication of currents directly affected by changing the bath potassium concentration (*K*_*bath*_). (B) Dependence of potassium (_*K*_) and leak (_*Leak*_) reversal potentials on *K*_*bath*_. Black dots indicate experimentally measured values for _*K*_ and _*Leak*_ from ***Koizumi and Smith (2008***). (C) Dependence of intrinsic cellular dynamics on *K*_*bath*_, *g*_*Leak*_ and *g*_*NaP*_. Black curve represents the relationship between *K*_*Bath*_ and *g*_*Leak*_ used in the full preBötC network. (D) Schematic diagram of size and connectivity probabilities of the rhythm and pattern generating populations within the preBötC model. (E) 2D plot between *g*_*NaP*_ and *g*_*Leak*_ showing the location of the intrinsic bursting regime for varied concentrations of *K*_*Bath*_. The distributions of neuronal conductances in the rhythm and the pattern generating populations are indicated by the blue dots and red squares, respectively.

To construct a model preBötC network, we linked rhythm and pattern generating subpopulations via excitatory synaptic connections within and between the two populations (Fig. 3D). We distinguished the two populations by endowing them with distinct distributions of persistent sodium current conductance (*g*_*NaP*_), as documented experimentally (***Del Negro et al., 2002a***; ***Koizumi and Smith, 2008***). In both populations, we maintained the dependence of *g*_*Leak*_ on *K*_*bath*_ (see Fig. 3C and E).

For the full preBötC network model, we first characterized the impact of changes in *K*_*bath*_ on network behavior without calcium dynamics by setting *P*_*SynCa*_ = 0. This network condition is analogous to *in vitro* preparations where all *Ca*^2+^ currents are blocked by *Cd*^2+^ and the preBötC can only generate burstlets (***Kam et al., 2013a***; ***Sun et al., 2019***). Not surprisingly, with calcium dynamics blocked, we found that the network can only generate small amplitude network oscillations (burstlets) that first emerge at approximately *K*_*bath*_ = 5 *mM* (Fig. 4A). Moreover, under these conditions, increasing *K*_*bath*_ results in an increase in the burstlet frequency and amplitude (Fig. 4B & C), which is consistent with experimental observations (***Kallurkar et al., 2020***).

**Figure 4.**
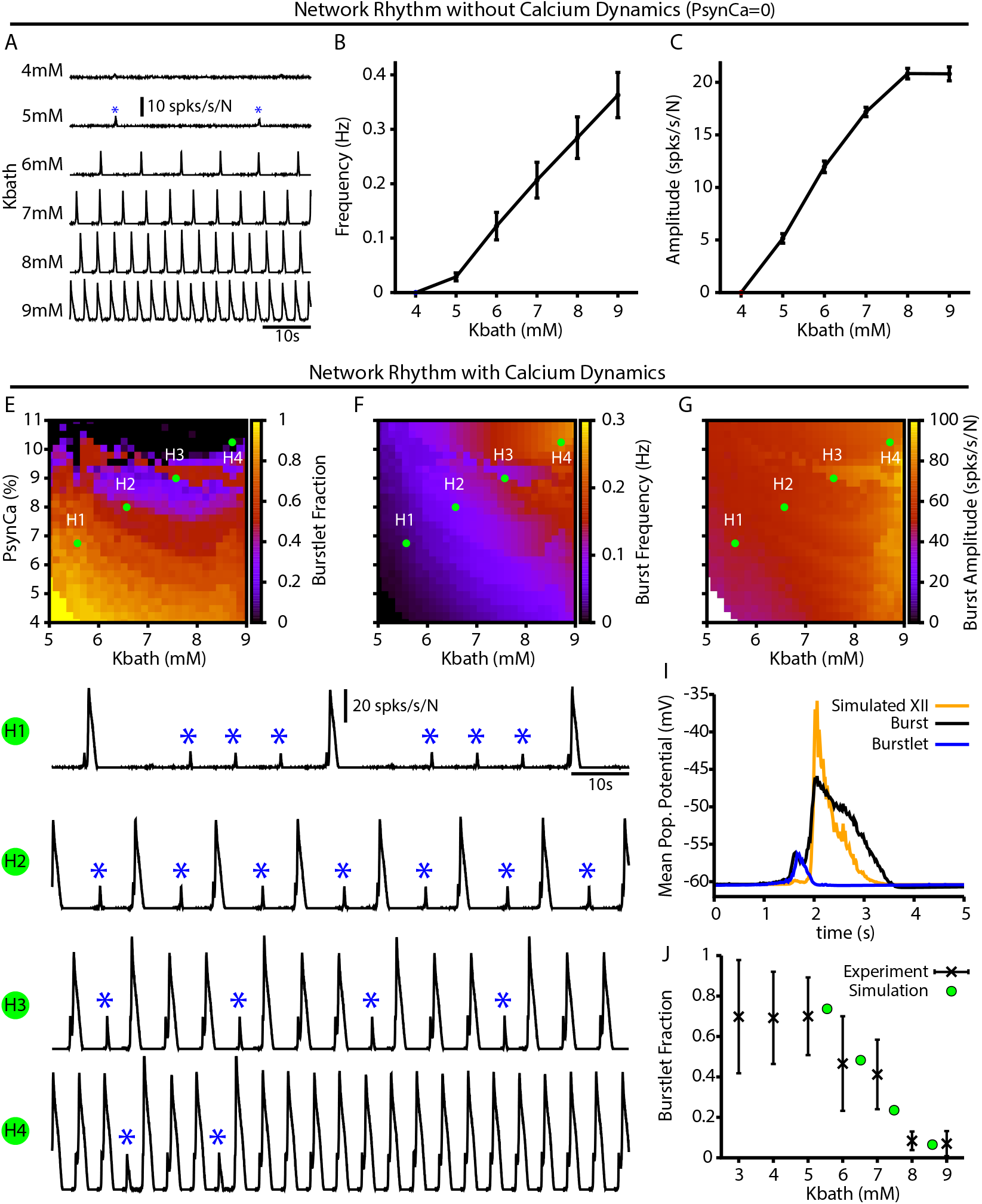
Burstlets and bursts in a 400 neuron preBötC network model with and without calcium dynamics (A) Rhythmogenic output of the simulated network without calcium dynamics (*P*_*SynCa*_ = 0) as a function of *K*_*Bath*_. These oscillations are considered burstlets as they are incapable of recruiting the pattern generating population without calcium dynamics. (B) Frequency and (C) amplitude of the burstlet oscillations as a function of *K*_*bath*_. (E-G) 2D plots characterizing the (E) burstlet fraction, (F) the burst frequency, and (G) the burst amplitude as a function of *K*_*bath*_ and *P*_*SynCa*_ (note that the *P*_*SynCa*_ range shown does not start at 0). (H1-H4) Example traces illustrating a range of possible burstlet fractions generated by the network. Burstlets are indicated by asterisks. (I) Overlay of the average population voltage during burst and burstlets. The hypoglossal output is calculated by passing the mean population through a sigmoid function *f* = −60.5 + 60/[1 + *e*^−(*x*+45)/2.5^]. (J) Burstlet fraction as a function of *K*_*bath*_ for the four example traces indicated in panels A-C. Experimental data is adapted from ***Kallurkar et al. (2020***).

In the full network with calcium dynamics (*P*_*SynCa*_ *>* 0), burstlets generated by the rhythmogenic subpopulation will trigger postsynaptic calcium transients in the pattern generating subpopulation. Therefore, in this set of simulations the burstlet activity of the rhythm generating population plays an analogous role to the square wave *Ca*^2+^ current in Fig. 1 and to bursts of the intrinsically rhythmic neuron in Fig. 2. In this case, the frequency of the postsynaptic *Ca*^2+^ oscillation is again controlled by *K*_*bath*_ and the *Ca*^2+^ amplitude is determined by the burstlet amplitude and *P*_*SynCa*_. Therefore, for this network, we characterized the burstlet fraction, burst frequency and burst amplitude – with a burst defined as an event in which a burstlet from the rhythm generating population recruits a burst in the pattern generating population – in the full preBötC model network as a function of *K*_*bath*_ and *P*_*SynCa*_ (Fig. 4E-G). We found that increasing *P*_*SynCa*_ or *K*_*bath*_ generally decreases the burstlet fraction, increases burst frequency, and slightly increases the burst amplitude (Fig. 4E-G and H1-H4). The decrease in the burstlet fraction with increasing *K*_*bath*_ or *P*_*SynCa*_ is caused by the increase in the burstlet amplitude (Fig. 4C) or in *Ca*^2+^ influx with each burstlet, respectively, both of which increase the *Ca*^2+^ transient in the pattern generating subpopulation. The increase in burst frequency with increases in *K*_*bath*_ or *P*_*SynCa*_ is due to the decreased burstlet fraction (i.e., the burstlet to burst transitions occurs on a greater proportion of cycles) and, in the case of *K*_*bath*_, by an increase in the burstlet frequency (Fig. 4B). The slight increase in burst amplitude with increasing *K*_*bath*_ is largely due to the increase in the burstlet amplitude (Fig. 4 C). Fig. 4I highlights the relative shape of burstlets and bursts as well as the delay between burstlet generation and recruitment of the pattern generating population and simulated hypoglossal output. Experimentally, it is likely that postsynaptic *Ca*^2+^ transients will increase with increasing *K*_*bath*_ due to the change in the resting *V*_*m*_ in non-rhythmic preBötC neurons (***Tryba et al., 2003***) relative to the voltage-gated activation dynamics of post-synaptic calcium channels (***Elsen and Ramirez, 1998***); see *Discussion* for a full analysis of this point. Interestingly, in our simulations, increasing *P*_*SynCa*_ (i.e. the amplitude of the postsynaptic calcium transients) with *K*_*bath*_ (Fig. 4 traces H1-H4) generated *K*_*bath*_-dependent changes in the burstlet fraction that are consistent with experimental observations (***Kallurkar et al., 2020***); see Fig. 4J.

Note that our model includes synaptic connections from pattern generating neurons back to rhythm generating neurons. These connections prolong activity of rhythmic neurons in bursts, relative to burstlets, which in turn yields a longer pause before the next event (e.g., Fig. 4H1). This effect can constrain event frequencies somewhat in the fully coupled network relative to the feed-forward case (e.g., frequencies in Fig. 4B exceed those in Fig. 4F for comparable *K*_*bath*_ levels).

### Calcium and *I*_*CAN*_ block have distinct effects on the burstlet fraction

Next, we further characterized the calcium dependence of the burstlet to burst transition in our model by simulating calcium blockade or *I*_*CAN*_ blockade by a progressive reduction of *P*_*SynCa*_ or *g*_*CAN*_, respectively. We found that complete block of synaptically triggered *Ca*^2+^ transients or *I*_*CAN*_ block eliminates bursting without affecting the underlying burstlet rhythm (Fig.5 A,B). Interestingly, progressive blockades of each of these two mechanisms have distinct effects on the burstlet fraction: blocking postsynaptic *Ca*^2+^ transients increases the burstlet fraction by increasing the number of burstlets required to trigger a network burst, whereas *I*_*CAN*_ block only slightly increases the burstlet fraction (Fig. 5C). In both cases, however, progressive blockade smoothly decreases the amplitude of network bursts, (Fig. 5D). The decrease in amplitude in the case of *I*_*CAN*_ block is due to derecruitment of neurons from the pattern forming subpopulation and a decrease in the firing rate of the neurons that remain active, whereas in the case of *Ca*^2+^ block the decrease in amplitude results primarily from derecruitment (Fig. 5E & F). These simulations provide mechanism-specific predictions that can be experimentally tested.

**Figure 5.**
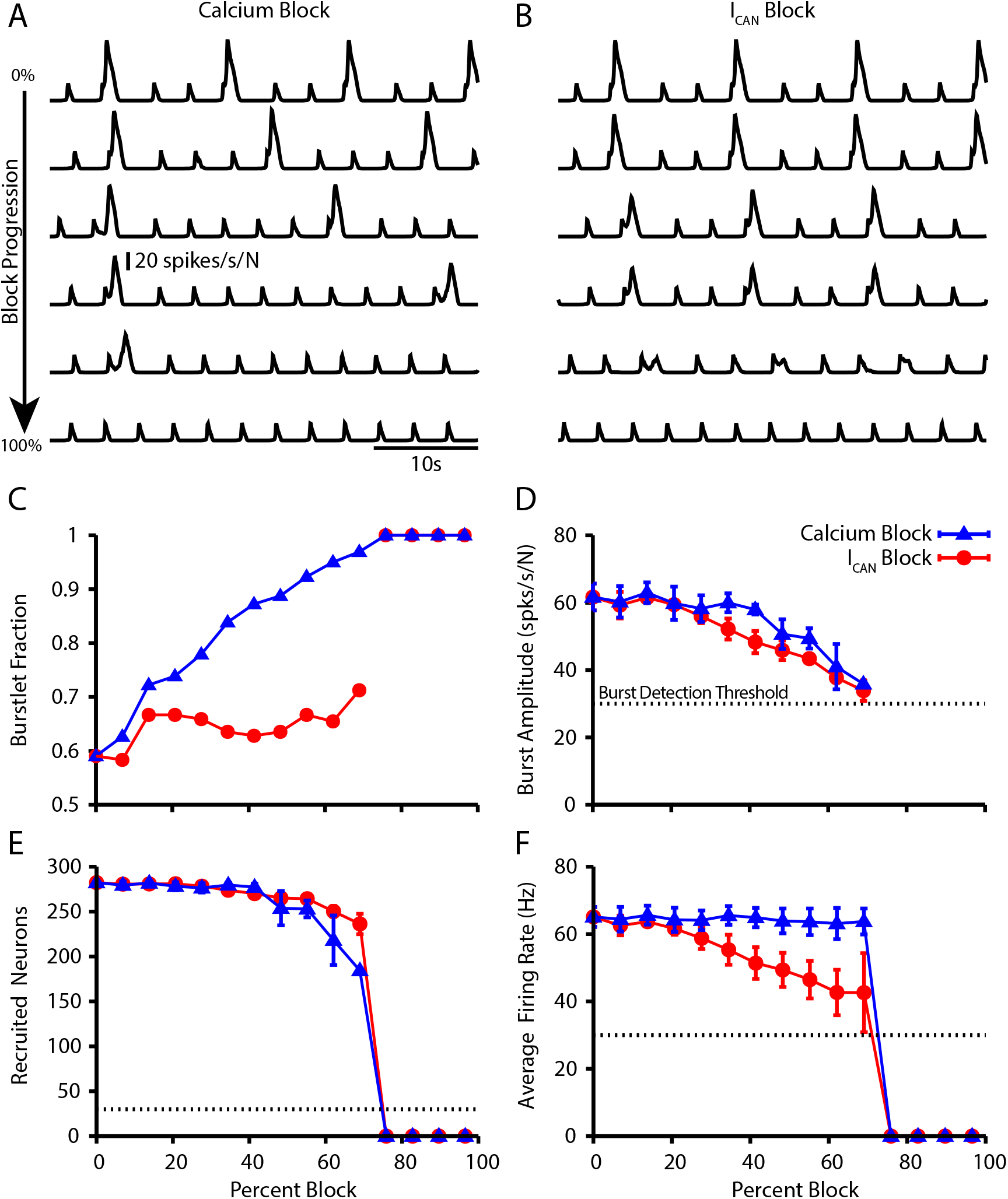
Effect of Ca^2+^ and CAN current blockade on burstlets and bursts. Network traces showing the affect of (A) Calcium current blockade (*P*_*SynCa*_ reduction) and (B) CAN current blockade (*g*_*CAN*_ reduction) on the period and amplitude of bursts. Effects of calcium or *I*_*CAN*_ blockade on (C) the burstlet fraction, (D) the amplitude of bursts and (E) the number of recruited and (F) peak firing rate of recruited neurons in pattern generating subpopulation during network bursts as a function of the blockade percentage.

### Dose dependent effects of opioids on the burstlet fraction

Recent experimental results by ***Baertsch et al. (2021***) showed that opioid application locally within the preBötC decreases burst frequency but also increases the burstlet fraction. In the preBötC, opioids affect neuronal dynamics by binding to the *µ*-opioid receptor (*µ*OR). The exact number of preBötC neurons expressing *µ*OR is unclear; however, the number appears to be small, with estimates ranging from 8 − 50% (***Bachmutsky et al., 2020***; ***Baertsch et al., 2021***; ***Kallurkar et al., 2021***).

Additionally, *µ*OR is likely to be selectively expressed on neurons involved in rhythm generation, given that opioid application in the preBötC primarily impacts burst frequency rather than amplitude (***Sun et al., 2019***; ***Baertsch et al., 2021***). Importantly, within the preBötC, opioids ultimately impact network dynamics through two distinct mechanisms: (1) hyperpolarization, presumably via activation of a G protein-gated inwardly rectifying potassium (GIRK) channel, and (2) decreased excitatory synaptic transmission, presumably via decreased presynaptic release (***Baertsch et al., 2021***).

Taking these considerations into account, we tested if our model could explain the experimental observations. Specifically, we simulated opioids as having a direct impact only on the neurons within the rhythmogenic population and their synaptic outputs (Fig. 6A). To understand how pre-BötC network dynamics are impacted by the two mechanisms though which opioids have been shown to act, we ran separate simulations featuring either activation of GIRK channels or block of the synaptic output from the rhythmogenic subpopulation (Fig. 6B-F). We found that both GIRK activation and synaptic block reduced the burst frequency (Fig. 6D) and slightly increased burst amplitude (Fig. 6E). The decreased frequency with synaptic block comes from an increase in the burstlet fraction, whereas GIRK activation kept the burstlet fraction constant while reducing the burstlet frequency (Fig. 6F). Finally, combining these effects, we observed that simultaneously increasing the GIRK channel conductance and blocking the synaptic output of *µ*OR-expressing neurons in our simulations generates slowing of the burst frequency and an increase in the burstlet fraction consistent with *in vitro* experimental data (Fig. 6D-G).

**Figure 6.**
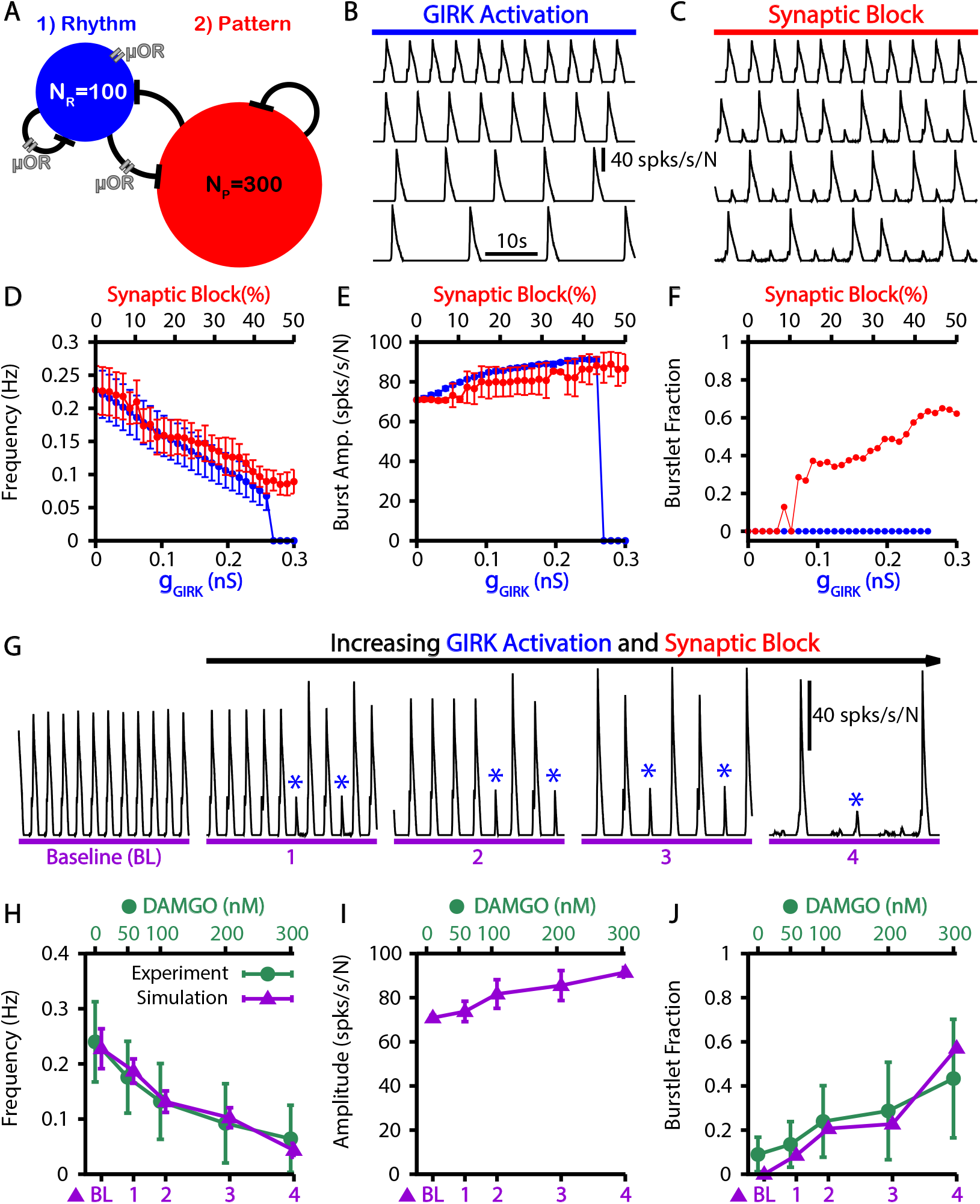
Simulated *µ*-opioid receptor (*µ*OR) activation by local DAMGO application in the preBötC and comparison with experimental data. (A) Schematic preBötC network diagram showing the location of *µ*OR. Example traces showing the effect of (B) *g*_*GIRK*_ channel activation and (C) synaptic block on the the network output. Quantification of *g*_*GIRK*_ activation or synaptic block by *µ*OR on the (D) burst frequency, (E) burst amplitude and (F) burstlet fraction. Error bars indicate SD. (G) Example traces showing the affects of progressive increases in *g*_*GIRK*_ and synaptic block on network output. Burstlets are indicated by blue asterisks. The parameters for each case are as follows: (BL) *g*_*GIRK*_ = 0.0 *nS*, _*γµ OR*_ = 0.0; (1) *g*_*GIRK*_ = 0.031034 *nS*, _*µ R*_ = 0.81034; (2) *g*_*GIRK*_ = 0.093103 *nS*,_*γ µ OR*_ = 0.7069; (3) *g*_*GIRK*_ = 0.14483 *nS*, _*γµ OR*_ = 0.68966; (4) *g*_*GIRK*_ = 0.19655 *nS*, _*γµ OR*_ = 0.58621. Comparison of experimental data and the affects of progressive increases in *g*_*GIRK*_ and synaptic block on the (H) frequency and (I) amplitude of bursts as well as (J) the burstlet fractions for the traces shown in (G). Experimental data (H, J) was adapted from ***Baertsch et al. (2021***); however, the changes in amplitude (I) were not quantified.

### Simultaneous stimulation of subsets of preBötC neurons elicits network bursts with long delays

Simultaneous stimulation of 4-9 preBötC neurons in *in vitro* slice preparations has been show to be sufficient to elicit network bursts with similar patterns to those generated endogenously (***Kam et al., 2013b***). These elicited bursts occur with delays of several hundred milliseconds relative to the stimulation time, which is longer than would be expected from existing models. Interestingly, in the current model, due to the dynamics of CICR, there is a natural delay between the onset of burstlets and the recruitment of the follower population that underlies the transition to a burst. Therefore, we investigated if our model could match and explain the observations seen in ***Kam et al. (2013b***).

In our model, we first calibrated our stimulation to induce a pattern of spiking that is comparable to the patterns generated in (***Kam et al., 2013b***) (10-15 spikes with decrementing frequency, Fig. 7A). We found that stimulation of 3-9 randomly selected neurons could elicit network bursts with delays on the order of hundreds of milliseconds (Fig. 7B & C). Next we characterized (1) the probability of eliciting a burst, (2) the delay in the onset of elicted bursts, and (3) the variability in delay, each as a function of the time of stimulation relative to the end of an endogenous burst (i.e., a burst that occurs without stimulation) and of the number of neurons stimulated (Fig. 7D-F). In general we found that increasing the number of stimulated neurons increases the probability of eliciting a burst and decreases the delay between stimulation and burst onset. Moreover, the probability of elicting a burst increases and the delay decreases as the time after an endogenous burst increases (Fig. 7G,H). Additionally, with its baseline parameter tuning, our model had a refractory period of approximately 1 *s* following an endogenous burst during which stimulation could not evoke a burst (Fig. 7). The refractory period in our model is longer than measured experimentally (500 *ms*) (***Kam et al., 2013b***).

**Figure 7.**
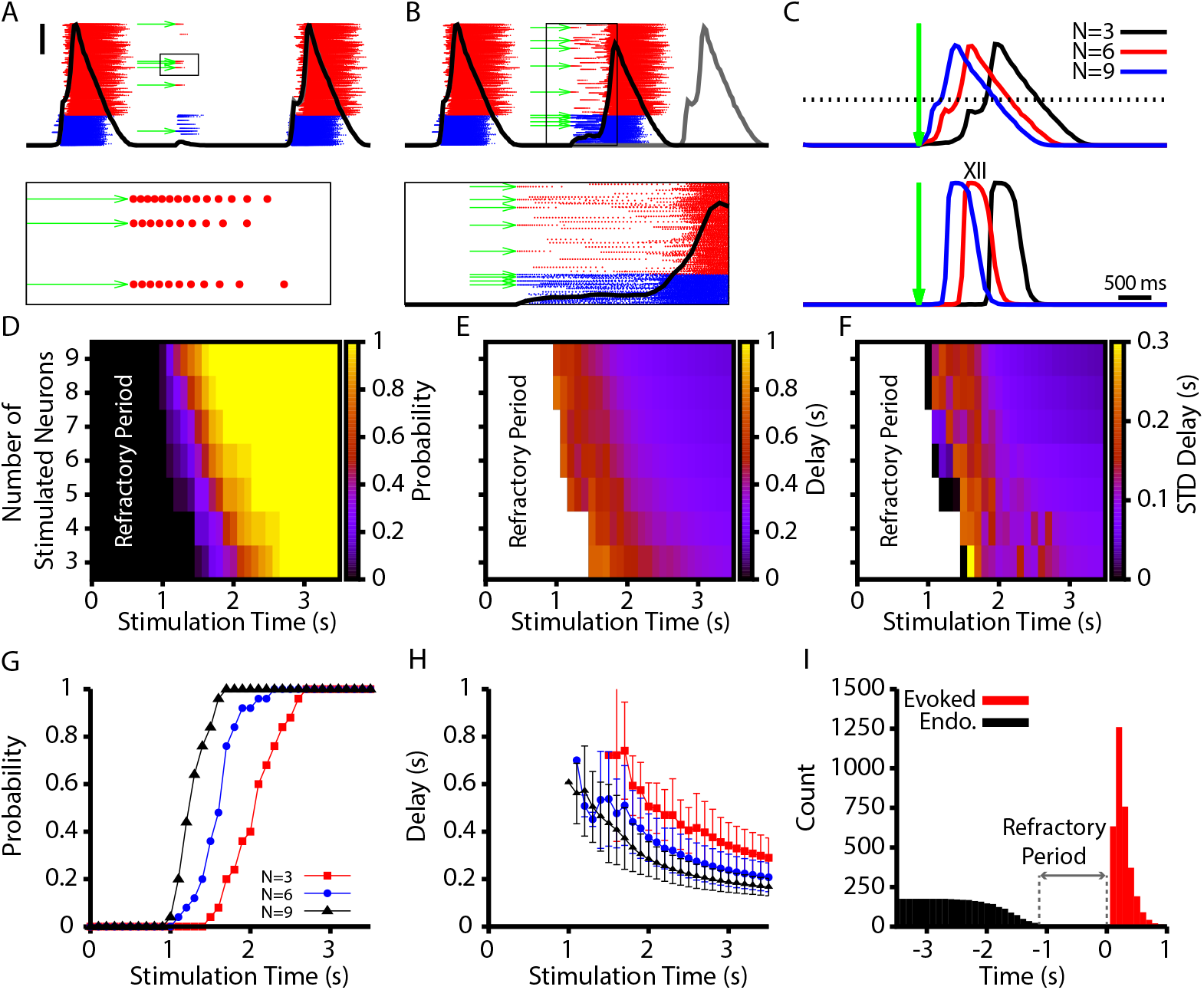
Evoked population bursts by simulated holographic stimulation of 3-9 preBötC neurons. (A) Raster plot of neuronal spiking triggered by simulated holographic stimulation of 6 preBötC neurons shortly after an endogenous burst and resulting failure to evoke a network burst. Black line represents the integrated population activity. Scale bar indicates 20 spikes/s/N. (Bottom panel) shows the spiking activity triggered in individual neurons by the simulated holographic stimulation. Panel duration is 1 *s*. (B) Example simulation where stimulation of 9 preBötC neurons evokes a network burst. Gray curve indicates timing of the next network burst if the network was not stimulated. (Bottom panel) Expanded view of the percolation process that is triggered by holographic stimulation on a successful trial. Panel duration is 1.75 *s*. (C) Example traces showing the delay between the stimulation time and the evoked bursts as a function of the number of neurons stimulated for the (top) integrated preBötC spiking and (bottom) simulated hypoglossal activity. (D-F) Characterization of (D) the probability of evoking a burst, (E) the mean delay of evoked bursts, and (F) the standard deviation of the delay as a function of the time after an endogenous burst and the number of neurons stimulated. (G) Probability and (H) delay as a function of the stimulation time for stimulation of 3, 6 or 9 neurons. Error bars in (H) indicate SD. (I) Histogram of evoked and endogenous bursts relative to the time of stimulation (*t* = 0 *s*) for all successful trials in all simulations; notice a 1 *s* refractory period.

To conclude our investigation, we examined how changes in the connection probability within the pattern forming population (*P*_*P P*_) affect the refractory period, probability, and delay of evoked bursts following simultaneous stimulation of 3-9 randomly selected neurons in the preBötC population. We focused on the pattern forming population because it comprises 75% of the preBötC population and, therefore, neurons from this population are most likely to be stimulated and the synaptic projections from these neurons are most likely to impact the properties of evoked bursts. These simulations were conducted with fixed network synaptic strength, defined as *S* = *N*_*P*_ · *P*_*P P*_ · *W*_*P P*_, where *W*_*P P*_ is adjusted to compensate for changes in *P*_*P P*_ to keep *S* constant.

With this scaling, we found that decreasing/increasing *P*_*P P*_ decreased/increased the refractory period (Fig. 8A-C) by impacting the probability of eliciting a burst in the period immediately after an endogenous burst (Fig. 8D-E). More specifically, the change in the probability of evoking a burst, with decreased/increased *P*_*P P*_, is indicated by a leftward/rightward shift in the probability vs stimulation time curves relative to a control level of *P*_*P P*_ (*P*_*P P*_ = 2%); see Fig. 8D,E. That is, relatively small connection probabilities with large connection strengths lead to network dynamics with a shorter refractory period when stimulation cannot elicit a burst and a higher probability that a stimulation at a fixed time since the last burst will evoke a new burst.

**Figure 8.**
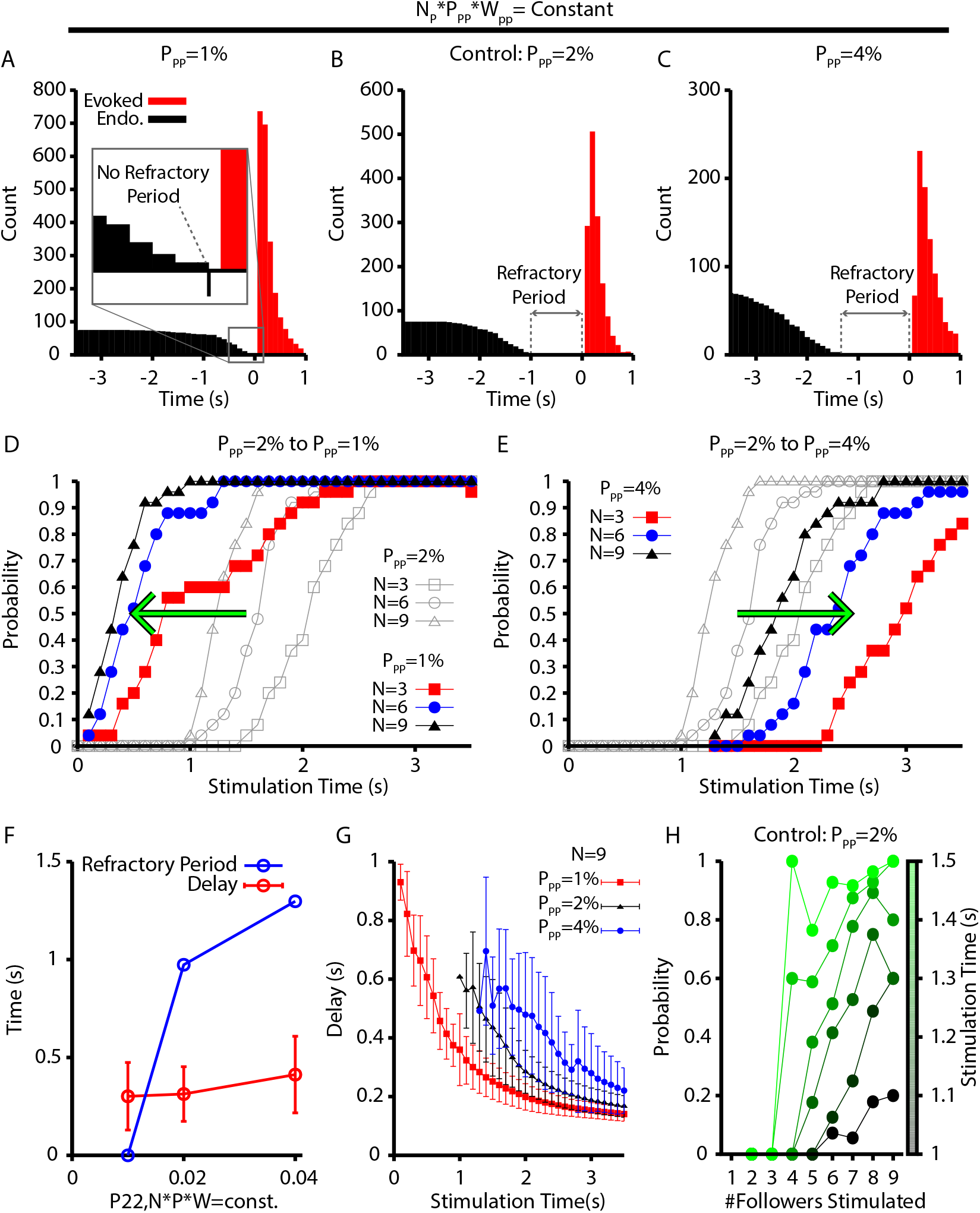
Refractory period and delay of evoked bursts following simulated holographic stimulation depends on the follower network connectivity. (A-C) Histogram of evoked and endogenous bursts relative to the time of stimulation (*t* = 0 *s*) for all successful trials where 3, 6 and 9 neurons were stimulated and for different connection probabilities (but fixed total network synaptic strength; i.e., *N*_*P*_ · *P*_*P P*_ · *W*_*P P*_ = *constant*) in the follower population: (A) *P*_*P P*_ = 1%; (B) *P*_*P P*_ = 2%; and (C) *P*_*P P*_ = 4%. (D & E) Effect of (D) decreasing (2% → 1%) and (E) increasing (2% → 4%) the connection probability in the follower population, *P*_*P P*_. (F) Refractory period and delay from stimulation to burst as functions of the connection probability for the simulations shown in A-E, with *N* · *P* · *W* = *const*. Error bars indicate SD. Notice that the refractory period increases with increasing connection probability. (G) Effect of *P*_*P P*_ on the delay to evoked bursts. (H) Probability of evoking a burst as a function of time (colorbar) and the number out of 9 stimulated neurons that are follower neurons for the baseline case of 2% connection probability.

It may seem surprising that networks with smaller connection probabilities exhibit a faster emergence of bursting despite their larger connection weights, since intuitively, with lower connection probabilities, fewer neurons could be recruited by each action potential, resulting in longer, more time-consuming activation pathways. A key point, however, is that when connection weights are larger, fewer temporally overlapping inputs are needed to recruit each inactive neuron. For example, fix *N*_*P*_ and *W*_*P P*_, take *P*_*P P*_ to scale as 1/*N*_*P*_, and assume that with this *W*_*P P*_, at least *r* inputs from active neurons are needed to activate an inactive neuron. We can approximate the expected number of neurons receiving *r* or more inputs from *A* active neurons by the expected number receiving *r* inputs, given by

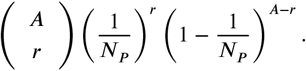

If we double *P*_*P P*_, halve *W*_*P P*_, and assume that now at least 2*r* inputs are needed for activation, then the corresponding approximation becomes

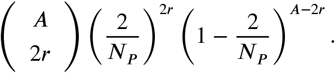

This is a smaller quantity than the first one for relevant parameter values (such as *N*_*P*_ = 300 and small *r* as indicated by the stimulation experiments), showing that increasing *P*_*P P*_ and proportionally scaling down *W*_*P P*_ reduces the chance of successful recruitment of inactive neurons by active neurons.

Interestingly, our simulations suggest that the connection probability in the pattern generating population must be between 1% and 2% to match the approximately 500 *ms* refractory period measured experimentally (***Kam et al., 2013b***) (Fig. 8F). Surprisingly, the mean and distribution of delays from stimulation to burst for all successfully elicited bursts are not strongly affected by changes in *P*_*P P*_ (Fig. 8F). For a given stimulation time and number of neurons stimulated, however, decreasing *P*_*P P*_ decreases the delay of elicited bursts (Fig. 8G). Finally, because the neurons in the pattern generating population appear to play a dominant role in determining if stimulation will elicit a network burst, we characterized how the number of pattern generating neurons stimulated, out of a total set of 9 stimulated neurons, affects the probability of eliciting a network burst as a function of stimulation time (Fig. 8H). These simulations were carried out under a baseline condition of *P*_*P P*_ = 2%. In general, we found that stimulating a relatively larger proportion of pattern generating neurons increased the probability of eliciting a network burst for all times after the approximately 1 *s* refractory period, as indicated by the positive slope in Fig. 8H.

## Discussion

Recent experiments have revealed a decoupling of respiratory rhythm generation and output patterning in the preBötC, which has given rise to the conceptual framework of burstlet theory. To date, however, this theory lacks the quantitative basis, grounded in underlying biophysical mechanisms, needed for its objective evaluation. To address this critical gap, in this computational study we developed a data-constrained biophysical model of the preBötC that generates burstlets and bursts as proposed by burstlet theory, with a range of features that match experimental observations. To summarize, we first show that calcium induced calcium release (CICR) from intracellular stores is a natural mechanism to periodically amplify postsynaptic calcium transients needed for *I*_*CAN*_ activation and recruitment of pattern-forming neurons into network bursts (Fig. 1). Next, we demonstrate that in a two-neuron network, CICR can convert baseline rhythmic activity into a mixture of bursts and burstlets, where the burstlet fraction depends largely on the magnitude of postsynaptic calcium transients (Fig. 2). In a larger preBötC network containing rhythm- and pattern-forming subpopulations with experimentally constrained intrinsic properties, population sizes and synaptic connectivity probabilities (Fig. 3), similar but more realistic activity patterns arise (Fig. 4). Moreover, we show that this model can match all of the key experimental underpinnings of burstlet theory, including the dependence of the burstlet fraction on extracellular potassium concentration (Fig. 4 I), the *Ca*^2+^ dependence of the burstlet-to-burst transition (Fig. 5), the effects of opioids on burst frequency and burstlet fraction (Fig. 6), and the long delay and refractory period of bursts evoked by holographic photostimulation of small subsets of preBötC neurons (Fig. 7 & 8).

### Insights into the mechanisms of burst (pattern) and burstlet (rhythm) generation in the inspiratory preBötC

Burstlet theory to date has largely been an empirical description of the observed features of bursts and burstlets. One idea that has been suggested is that rhythm generation is driven by a stochastic percolation process in which tonic spiking across the rhythm-generating population gradually synchronizes during the inter-burst-interval to generate the burstlet rhythm. Subsequently, a burst (i.e. motor output) only occurs if the burstlet is of sufficient magnitude, resulting from sufficient synchrony, to trigger all-or-none recruitment of the pattern-forming subpopulation (***Kam et al., 2013a,b***; ***Feldman and Kam, 2015***; ***Cui et al., 2016***; ***Kallurkar et al., 2020***; ***Ashhad and Feldman, 2020***). This theory, however, does not identify or propose specific biophysical mechanisms capable of generating a quantitative explanation of the underlying cellular and network level dynamics, fails to capture the *Ca*^2+^ dependence of the burst-to-burstlet transition, and cannot explain how extracellular potassium concentration impacts the burstlet fraction. Our simulations support an alternative view that builds directly from previous computational studies (***Jasinski et al., 2013***; ***Phillips et al., 2019***; ***Phillips and Rubin, 2019***; ***Phillips et al., 2021***), which robustly reproduce a wide array of experimental observations. Specifically, in this study we show that amplification of postsynaptic calcium transients in the pattern-generating subpopulation (triggered by burstlets) provides a natural mechanism capable of explaining the *Ca*^2+^ dependence of the burstlet-to-burst transition.

Importantly, we find that the burstlet fraction is determined by the probability that a burstlet will trigger CICR in the pattern forming subpopulation. In the model, this probability is determined by the magnitude of postsynaptic calcium transients as well as the activation dynamics of the IP3 receptor and the SERCA pump. Therefore, to explain the decrease in the burstlet fraction with increasing extracellular *K*_*bath*_, the magnitude of the burstlet-triggered postsynaptic calcium transients must increase with *K*_*bath*_. Some of this rise can result directly from the increase in burstlet amplitude with increasing *K*_*bath*_ (see (***Kallurkar et al., 2020***) and Fig. 4 C). To fully match the experimentally observed relationship between *K*_*bath*_ and the burstlet fraction (Fig. 4 J), we also explicitly increased the parameter *P*_*SynCa*_, which sets the proportion of the postsynaptic calcium current carried by *Ca*^2+^. Thus, our model predicts that the magnitude of postsynaptic *Ca*^2+^ transients triggered by EPSPs should increase as *K*_*bath*_ is elevated.

This same prediction arises from considering the voltage-dependent properties of *Ca*^2+^ channels characterized in preBötC neurons and the changes in the membrane potential of non-rhythmogenic (i.e. pattern-forming) neurons as a function of *K*_*bath*_. Specifically, it is likely that voltage-gated calcium channels are involved in generating the postsynaptic *Ca*^2+^ transients, as dendritic *Ca*^2+^ transients have been shown to precede inspiratory bursts and to be sensitive to *Cd*^2+^, a calcium channel blocker (***Del Negro et al., 2011***). Consistent with this idea, *Cd*^2+^-sensitive *Ca*^2+^ channels in preBötC neurons appear to be primarily localized in distal dendritic compartments (***Phillips et al., 2018***). Voltage-gated calcium channels in the preBötC start to activate at approximately −65 *mV* (***Elsen and Ramirez, 1998***) and importantly, the mean somatic resting membrane potential of non-rhythmogentic preBötC neurons increases from −67.034 *mV* to −61.78 *mV* when extracellular potassium concentration is elevated from 3 *mM* to 8 *mM* (***Tryba et al., 2003***). Moreover, at *K*_*Bath*_ = 9 *mM*, EPSPs in the preBötC are on the order of 2 − 5 *mV* (***Kottick and Del Negro, 2015***; ***Morgado-Valle et al., 2015***; ***Baertsch et al., 2021***) and the amplitude of EPSCs has been shown to decrease as *K*_*bath*_ is lowered (***Okada et al., 2005***). Putting together these data on resting membrane potential and EPSP sizes, we deduce that when *K*_*Bath*_ = 3 *mM*, the magnitude of EPSPs may not reach voltages sufficient for significant activation of voltage-gated *Ca*^2+^ channels. As *K*_*bath*_ is increased, however, increases in the membrane potential of pattern-forming neurons and EPSP magnitude are predicted to increase the magnitude of EPSPs triggered by postsynaptic calcium transients. This is exactly the effect that is captured in the model by an increase in *P*_*SynCa*_.

The idea that dendritic post-synaptic *Ca*^2+^ transients and *I*_*CAN*_ activation play a critical role in regulating the pattern of preBötC dynamics is well supported by experimental and computational studies. Specifically, the dendritic *Ca*^2+^ transients that precede inspiratory bursts (***Del Negro et al., 2011***) have been shown to travel in a wave to the soma, where they activate TRPM4 currents (*I*_*CAN*_) (***Mironov, 2008***). Moreover, the rhythmic depolarization of otherwise non-rhythmogenic neurons (inspiratory drive potential) depends on *I*_*CAN*_ (***Pace et al., 2007a***), while the inspiratory drive potential is not dependent on *Ca*^2+^ transients driven by voltage-gated calcium channels expressed in the soma (***Morgado-Valle et al., 2008***). Finally, pharmacological blockade of TRPM4 channels, thought to represent the molecular correlates of *I*_*CAN*_, reduces the amplitude of preBötC motor output without impacting the rhythm. These experimental findings were incorporated into and robustly reproduced in a recent computational model (***Phillips et al., 2019***). Consistent with these findings, this previous model suggests that rhythm generation arises from a small subset of preBötC neurons, which form an *I*_*NaP*_ -dependent rhythmogenic kernel (i.e. burstlet rhythm generator), and that rhythmic synaptic drive from these neurons triggers post-synaptic calcium transients, *I*_*CAN*_ activation, and amplification of the inspiratory drive potential, which spurs bursting in the rest of the network. The current study builds on this previous model by explicitly defining rhythm- and pattern-generating neuronal subpopulations (see Fig. 3) and by incorporating the mechanisms required for CICR and intermittent amplification of post-synaptic calcium transients.

Calcium-induced calcium release mediated by the SERCA pump and the IP3 receptor has long been suspected to be involved in the dynamics of preBötC rhythm and/or pattern generation (***Pace et al., 2007a***; ***Crowder et al., 2007***; ***Mironov, 2008***; ***Toporikova et al., 2015***) and has been explored in individual neurons and network models of the preBötC (***Toporikova and Butera, 2011***; ***Jasinski et al., 2013***; ***Rubin et al., 2009***; ***Wang and Rubin, 2020***). Experimental studies have not clearly established the role of CICR from ER stores in respiratory circuits, however. For example, ***Mironov (2008***) showed that the transmission of calcium waves that travel from the dendrites to the soma is blocked by local application of thapsigargin, a SERCA pump inhibitor. In a separate study, however, block of the SERCA pump by bath application of thapsigargin (2 − 20 *µM*) or cyclopiazonic acid (CPA) (30 − 50 *µM*) did not significantly affect the amplitude or frequency of hypoglossal motor output in *in vitro* slice preparations containing the preBötC. It is possible that the negative results presented by the latter work occur due to the failure of pharmacological agents to fully penetrate the slice and diffuse across the cell membranes to reach their intracellular targets. Alternatively, the role of CICR may be dynamically regulated depending on the state of the preBötC network. For example the calcium concentration at which the IP3 receptor is activated is dynamically regulated by IP3 (***Kaftan et al., 1997***) and therefore, activity- or neuromodulatory-dependent changes in the cytoplasmic *Ca*^2+^ and/or IP3 concentration may impact ER *Ca*^2+^ uptake and release dynamics. Store operated *Ca*^2+^ dynamics are also impacted by the transient receptor potential canonical 3 (TRPC3) channels (***Salido et al., 2009***), which are expressed in the preBötC, and manipulation of TRPC3 has been shown to impacted burst amplitude and regularity (***Tryba et al., 2003***; ***Koizumi et al., 2018***) as would be predicted by this model. It is also possible that calcium release is mediated by the ryanodine receptor, an additional calcium activated channel located in the ER membrane (***Lanner et al., 2010***), since bath application of CPA (100 *µM*) and ryanodine (10 *µM*) removed large amplitude oscillations in recordings of preBötC population activity (***Toporikova et al., 2015***).

Finally, we note that while various markers can be used to define distinct subpopulations of neurons within the preBötC, our model cannot determine which of these ensembles are responsible for rhythm and pattern generation. Past experiments have examined the impact of optogenetic inhibition, applied at various intensities to subpopulations associated with specific markers, on the frequency of inspiratory neural activity, but this activity was measured by motor output, not within the preBötC itself (***Tan et al., 2008***; ***Cui et al., 2016***; ***Koizumi et al., 2016***). According to burstlet theory and our model, slowed output rhythmicity could derive from inhibition of rhythm-generating neurons, due to a reduced frequency of burstlets, and from inhibition of pattern-generating neurons, due to a reduced success rate of burst recruitment. Thus, measurements within the preBötC will be needed in order to assess the mapping between subpopulations of preBötC neurons and roles in burstlet and burst production.

### Additional comparisons to experimental results

In our model, a burstlet rhythm first emerges at a *K*_*bath*_ of approximately 5 *mM*, whereas in the experiments of ***Kallurkar et al. (2020***), the burstlet rhythm continues even down to 3 *mM*. To explain this discrepancy, we note that our model assumes that the extracellular potassium concentration throughout the network is equal to *K*_*bath*_. Respiratory circuits appear to have some buffering capacity, however, such that for *K*_*bath*_ concentrations below approximately 5 *mM* the extracellular *K*^+^ concentration remains elevated above *K*_*bath*_ (***Okada et al., 2005***). The *K*_*bath*_ range over which our model generates a rhythm would extend to that seen experimentally if extracellular *K*^+^ buffering were accounted for. This buffering effect can also explain why the burstlet fraction remains constant in experimental studies when *K*_*bath*_ is lowered from 5 *mM* to 3 *mM* (***Kallurkar et al., 2020***). Our model also does not incorporate short-term extracellular potassium dynamics that may impact the ramping shape of burstlet onset (***Abdulla et al., 2021***).

Although our model incorporates various key biological features, it does not include some of the biophysical mechanisms that are known to shape preBötC patterned output or that are hypothesized to contribute to the properties described by burstlet theory. For example, the M-current associated with KCNQ potassium channels has been shown to impact burst duration by contributing to burst termination (***Revill et al., 2021***). Additionally, intrinsic conductances associated with a hyperpolarization-activated mixed cation current (*I*_*h*_) and a transient potassium current (*I*_*A*_) are hypothesized to be selectively expressed in the pattern- and rhythm-generating preBötC subpopulations (***Picardo et al., 2013***; ***Phillips et al., 2018***). Thus, our model predicts that while these currents may impact quantitative properties of burstlets and bursts, they are not critical for the presence of burstlets and their transformation into bursts. Finally, the current model does not include a population of inhibitory preBötC neurons. Inhibition is involved in regulating burst amplitude (***Baertsch et al., 2018***), but it does not have a clear role in burst or burstlet generation, and therefore inhibition was omitted from this work.

Importantly, our model does robustly reproduce all of the key experimental observations underlying burstlet theory. Not surprisingly, block of calcium transients or *I*_*CAN*_ in our model eliminates bursts without affecting the underlying rhythm (Fig. 5), which is consistent with experimental observations (***Kam et al., 2013b***; ***Sun et al., 2019***). Interestingly, our model also provides the experimentally testable predictions that blocking calcium transients will increase the burstlet fraction while *I*_*CAN*_ block will have no effect on this fraction, whereas both perturbations will smoothly reduce burst amplitude. Interestingly, the calcium-dependent mechanisms that we include in our model pattern-generating population have some common features with a previous model that suggested the possible existence of two distinct preBötC neuronal populations responsible for eupneic burst and sigh generation, respectively, which also included excitatory synaptic transmission from the former to the latter (***Toporikova et al., 2015***). In the eupnea-sigh model, however, the population responsible for low-frequency, high-amplitude sighs was capable of rhythmic burst generation even without synaptic drive, in contrast the pattern-generation population as tuned in our model. Also in contrast to the results on bursts considered in our study, sigh frequency in the earlier model did not vary with extracellular potassium concentration and sigh generation required a hyperpolarization-activated inward current, *I*_*h*_.

We also considered the effects of opioids in the context of burstlets and bursts, a topic that has not been extensively studied. It is well established that opioids slow the preBötC rhythm in *in vitro* slice preparations; however, the limited results presented to date on effects of opioids on the burstlet fraction are inconsistent. Specifically, ***Sun et al. (2019***) found that application of the *µ*-opioid receptor agonist DAMGO at 10 *nM* and 30 *nM* progressively decreased the preBötC network frequency but had no impact on the burstlet fraction before the network rhythm was eventually abolished at approximately 100 *nM*. Similarly, ***Baertsch et al. (2021***) found that DAMGO decreased the preBötC network frequency in a dose-dependent fashion; however, in these experiments the network was less sensitive to DAMGO, maintaining rhythmicity up to approximately 300 *nM*, and the burstlet fraction increased with increasing DAMGO concentration. The inconsistent effects of DAMGO on the burstlet fraction across these two studies can be explained by our simulations based on the different sensitivities of these two preparations to DAMGO and the two distinct mechanisms of action of DAMGO on neurons that express *µ*OR – decreases in excitability and decreases in synaptic output of neurons – identified by ***Baertsch et al. (2021***). In our simulations we show that the decreased excitability resulting from activation of a GIRK channel only impacts frequency, whereas decreasing the synaptic output of *µ*OR-expressing neurons results in an increase in the burstlet fraction and a decrease in burst frequency (Fig. 6). In experiments, suppression of synaptic output does not appear to occur until DAMGO concentrations are above approximately 50 *nM* (***Baertsch et al., 2021***). Therefore, it is not surprising that DAMGO application did not strongly impact the burstlet fraction before the rhythm was ultimately abolished in ***Sun et al. (2019***), due to the higher DAMGO sensitivity of that particular experimental preparation, as indicated by the lower dose needed for rhythm cessation.

### Mixed-mode oscillations

Mixed-mode oscillations, in which intrinsic dynamics of a nonlinear system naturally lead to alternations between small- and large-amplitude oscillations (***Del Negro et al., 2002c***; ***Bertram and Rubin, 2017***), are a mechanism that has been previously proposed to underlie bursts and burstlets, under the assumption of differences in intrinsic oscillation frequencies across preBötC neurons (***Bacak et al., 2016***). This mechanism was not needed to explain the generation of bursts and burstlets in the current model, however. Moreover, systems with mixed-mode oscillations can show a wide range of oscillation amplitudes under small changes in conditions and only emerge when *K*_*bath*_ elevated above 9 *mM* (***Del Negro et al., 2002c***). These properties are not consistent with the burst and burstlet amplitudes or *K*_*bath*_-dependent changes in the burstlet fraction seen experimentally (***Kallurkar et al., 2020***) and in our model.

### Holographic photostimulation, percolation and rhythm generation

Experimental data supporting burstlet theory has shown that burstlets are the rhythmogenic event in the preBötC. However, although burstlet theory is sometimes referenced as a theory of respiratory rhythm generation, the actual mechanisms of burstlet rhythm generation remain unclear. One idea that has been suggested is that rhythm generation is driven by a stochastic percolation process in which tonic spiking across the rhythm-generating population gradually synchronizes during the inter-burst-interval to generate the burstlet rhythm (***Ashhad and Feldman, 2020***; ***Slepukhin et al., 2020***). In this framework, a burst (i.e. motor output) only occurs if the burstlet is of sufficient magnitude, resulting from sufficient synchrony, to trigger all-or-none recruitment of the pattern-forming subpopulation (***Kam et al., 2013a,b***; ***Feldman and Kam, 2015***; ***Kallurkar et al., 2020***; ***Ashhad and Feldman, 2020***; ***Slepukhin et al., 2020***).

The idea that burstlets are the rhythmogenic event within the preBötC is supported by the observation that block of voltage-gated *Ca*^2+^ channels by *Cd*^2+^ eliminates bursts without affecting the underlying burstlet rhythm (***Kam et al., 2013a***; ***Sun et al., 2019***). However, the rhythmogenic mechanism based on percolation is speculative and comes from two experimental observations. The first is that the duration and slope (i.e., shape) of the burstlet onset are statistically indistinguishable from the ramping pre-inspiratory activity that immediately precedes inspiratory bursts (***Kallurkar et al., 2020***). We note, however, that this shape of pre-inspiratory activity can arise through intrinsic mechanisms at the individual neuron level (***Abdulla et al., 2021***). The second observation evoked in support of the percolation idea is that holographic photstimulaton of small subsets (4 − 9) of preBötC neurons can elicit bursts with delays lasting hundreds of milliseconds (***Kam et al., 2013b***). These delays are longer than could be explained with existing preBötC models and have approximately the same duration as the pre-inspiratory activity and burstlet onset hypothesized to underlie the rhythm. According to the percolation hypothesis of burstlet rhythm generation, these long delays result from the specific topological architecture of the preBötC, recently proposed to be a heavy-tailed synaptic weight distribution in the rhythmogenic preBötC subpopulation (***Slepukhin et al., 2020***).

Interestingly, the model presented here naturally captures the long delays characterized by ***Kam et al. (2013b***), and stimulation of small subsets of neurons triggers a growth in population activity in the lead up to a burst that could be described as percolation (Fig. 7B). Our model does not require a special synaptic weight distribution to generate the long delays, however. Indeed, our model suggests that the long delays between simulation and burst generation are due in large part to the dynamics of the pattern-forming population, as probabilistically these neurons are the most likely to be stimulated and they appear to play a dominant role in setting the timing of the elicited burst response (Fig. 8H). Moreover, the dynamics of this population is strongly impacted by the CICR mechanism proposed here, which is required for burst generation. Interestingly, to match the 500 *ms* refractory period following an endogenous burst during which holographic stimulation cannot elicit a burst, our model predicts that the connection probability in the pattern generating preBötC subpopulation must be between 1% and 2%, which is consistent with available experimental data (***Ashhad and Feldman, 2020***).

Thus, taken together, previous modeling and our work offer two alternative, seemingly viable hypotheses about the source of the delay between holographic stimulation and burst onset, each related to a proposed mechanism for burstlet and burst generation. Yet additional arguments call into question aspects of the percolation idea. If the burstlet rhythm is driven by a stochastic percolation process, then the period and amplitude of burstlets should be stochastic, irregular, and broadly distributed. Moreover, in the proposed framework of burstlet theory, the pattern of bursts and burstlets for a given burstlet fraction would also be stochastic, since the burstlet-to-burst transition is thought to be an all-or-none process that depends on the generation of a burstlet of sufficient magnitude. Example traces illustrating a mixture of bursts and burstlets typically show a pattern of multiple burstlets followed by a burst that appears to regularly repeat (***Kam et al., 2013b***; ***Sun et al., 2019***; ***Kallurkar et al., 2020***) and hypoglossal output timing has also been found to exhibit high regularity ***Kam et al. (2013b***), however, suggesting that the burstlet-to-burst transition is not dependent on the synchrony and hence magnitudes of individual burstlets but rather on a slow process that gradually evolves over multiple burstlets. The regularity and patterns of burstlets and bursts that arise from such a process in our model match well with those observed experimentally.

We note that the burstlet-to-burst transition mechanism proposed here, based on CICR from ER stores, depends on rhythmic inputs from the rhythm-generating to the pattern-generation population; however, it is independent of the mechanism of rhythm generation. In our simulations, rhythm generation depends on the slowly inactivating persistent sodium current (*I*_*NaP*_). The role of *I*_*NaP*_ in preBötC inspiratory rhythm generation is a contentious topic within the field, largely due to the inconsistent effects of *I*_*NaP*_ block. We chose to use *I*_*NaP*_ in as the rhythmogenic mechanism in the burstlet population for a number of reasons: (1) consideration of the pharmacological mechanism of action and nonuniform effects of drug penetration can explain the seemingly contradictory experimental findings relating to *I*_*NaP*_ (***Phillips and Rubin, 2019***), (2) *I*_*NaP*_ -dependent rhythm generation is a well-established and understood idea (***Butera et al., 1999***), (3) recent computational work on which the current model is based suggests that rhythm generation occurs in a small, *I*_*NaP*_ -dependent rhythmogenic kernel that is analogous to the burstlet population (***Phillips et al., 2019***), and predictions from this model that depend on the specific proposed roles of *I*_*NaP*_ and *I*_*CAN*_ in rhythm and pattern formation have been experimentally confirmed in a recent study (***Phillips et al., 2021***). It is important to note, however, that the findings about burstlets and bursts presented in this work would have been obtained if the burstlet rhythm was imposed (Fig. 1) or if burstlets were generated by some other means, such as by the percolation mechanism proposed by burstlet theory.

## Conclusions

This study has developed the first model-based description of the biophysical mechanism underlying the generation of bursts and burstlets in the inspiratory preBötC. As suggested by burstlet theory and other previous studies, rhythm and pattern generation in this work are represented by two distinct preBötC subpopulations. A key feature of our model is that generation of network bursts (i.e. motor output) requires amplification of postsynaptic *Ca*^2+^ transients by CICR in order to activate *I*_*CAN*_ and drive bursting in the rest of the network. Moreover, the burstlet fraction depends on rate of *Ca*^2+^ buildup in intracellular stores, which is impacted by *K*_*bath*_-dependent modulation of preBötC excitability. These ideas complement other recent findings on preBötC rhythm generation (***Phillips et al., 2019***; ***Phillips and Rubin, 2019***; ***Phillips et al., 2021***), together offering a unified explanation for a large body of experimental findings on preBötC inspiratory activity that offer a theoretical foundation on which future developments can build.

## Methods and Materials

### Neuron Model

Model preBötC neurons include a single compartment and incorporate Hodgkin-Huxley style conductances adapted from previously described models (***Jasinski et al., 2013***; ***Phillips et al., 2019***; ***Phillips and Rubin, 2019***) and/or experimental data as detailed below. The membrane potential of each neuron is governed by the following differential equation:

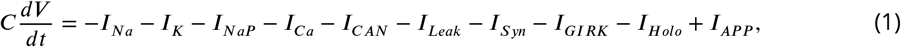

where *C* = 36 *pF* is the membrane capacitance and each *I*_*i*_ represents a current, with *i* denoting the current’s type. The currents include the action potential generating Na^+^ and delayed rectifying K^+^ currents (*I*_*Na*_ and *I*_*K*_), persistent Na^+^ current (*I*_*NaP*_), voltage-gated Ca^2+^ current (*I*_*Ca*_), Ca^2+^-activated non-selective cation (CAN) current (*I*_*CAN*_), K^+^ dominated leak current (*I*_*Leak*_), synaptic current (*I*_*Syn*_), *µ*-opioid receptor activated G protein-coupled inwardly-rectifying K^+^ current (*I*_*GIRK*_), and a holographic photostimulation current (*I*_*Holo*_). *I*_*AP P*_ denotes an applied current injected from an electrode. The currents are defined as follows:

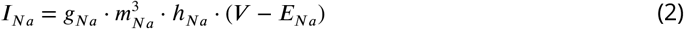

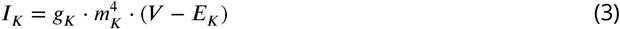

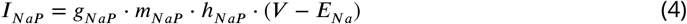

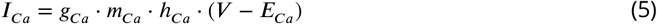

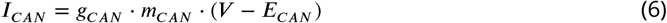

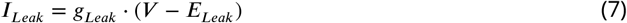

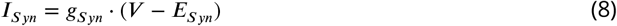

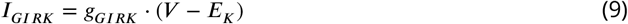

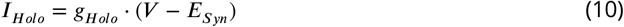

where *g*_*i*_ is the maximum conductance,*E* _*i*_ is the reversal potential, and *m*_*i*_ and *h*_*i*_ are gating variables for channel activation and inactivation for each current *I*_*i*_. The glutamatergic synaptic conductance *g*_*Syn*_ is dynamic and is defined below. The values used for the *g*_*i*_ and *E* _*i*_ appear in Table 1.

**Table 1.** Ionic Channel Parameters.

| Channel | Parameters |  |  |  |  |
| --- | --- | --- | --- | --- | --- |
| $I_{Na}$ | $g_{Na} = 150 \text{ nS}$<br>$m_{1/2} = -43.8 \text{ mV}$<br>$h_{1/2} = -67.5 \text{ mV}$ | $E_{Na} = 26.54 \cdot \ln(Na_{out}/Na_{in})$<br>$k_m = 6.0 \text{ mV}$<br>$k_h = -11.8 \text{ mV}$ | $Na_{in} = 15 \text{ mM}$<br>$\tau_{max}^m = 0.25 \text{ ms}$<br>$\tau_{max}^h = 8.46 \text{ ms}$ | $Na_{out} = 120 \text{ mM}$<br>$\tau_{1/2}^m = -43.8 \text{ mV}$<br>$\tau_{1/2}^h = -67.5 \text{ mV}$ | $k_{\tau}^m = 14.0 \text{ mV}$<br>$k_{\tau}^h = 12.8 \text{ mV}$ |
| $I_K$ | $g_K = 220 \text{ nS}$<br>$A_{\alpha} = 0.011$<br>$A_{\beta} = 0.17$ | $E_K = 26.54 \cdot \ln(K_{bath}/K_{in})$<br>$B_{\alpha} = 44.0 \text{ mV}$<br>$B_{\beta} = 49.0 \text{ mV}$ | $K_{in} = 125$<br>$k_{\alpha} = 5.0 \text{ mV}$<br>$k_{\beta} = 40.0 \text{ mV}$ | $K_{Bath} = V_{AR}$ | |
| $I_{NaP}$ | $g_{NaP} = N(\mu, \sigma)$, See Table 2<br>$m_{1/2} = -47.1 \text{ mV}$<br>$h_{1/2} = -60.0 \text{ mV}$ | $k_m = 3.1 \text{ mV}$<br>$k_h = -9.0 \text{ mV}$ | $\tau_{max}^m = 1.0 \text{ ms}$<br>$\tau_{max}^h = 5000 \text{ ms}$ | $\tau_{1/2}^m = -47.1 \text{ mV}$<br>$\tau_{1/2}^h = -60.0 \text{ mV}$ | $k_{\tau}^m = 6.2 \text{ mV}$<br>$k_{\tau}^h = 9.0 \text{ mV}$ |
| $I_{Ca}$ | $g_{Ca} = 0.0065 \text{ pS}$<br>$m_{1/2} = -27.5 \text{ mV}$<br>$h_{1/2} = -52.4 \text{ mV}$ | $E_{Ca} = 13.27 \cdot \ln(Ca_{out}/Ca_{in})$<br>$k_m = 5.7 \text{ mV}$<br>$k_h = -5.2 \text{ mV}$ | $\tau_m = 0.5 \text{ ms}$<br>$\tau_h = 18.0 \text{ ms}$ | $Ca_{out} = 4.0 \text{ mM}$ | |
| $I_{CAN}$ | $g_{CAN} = N(\mu, \sigma)$, See Table 2 | $E_{CAN} = 0.0 \text{ mV}$ | $Ca_{1/2} = 0.00074 \text{ mM}$ | $n = 0.97$ | |
| $I_{Leak}$ | $g_{Leak} = N(\mu, \sigma)$, See Table 2<br>$P_{Na} = 1$ | $E_{Leak} = -26.54 \cdot \ln[(P_{Na} * Na_{in} + P_K * K_{in}) / (P_{Na} * Na_{out} + P_K * K_{bath})]$<br>$P_K = 42$ | | | |
| $I_{Syn}$ | $g_{Syn} = V_{AR}$, See Eq. 25 | $E_{Syn} = 0.0 \text{ mV}$ | $\tau_{Syn} = 5.0 \text{ ms}$ | | |
| $I_{GIRK}$ | $g_{GIRK} = 0 - 0.3 \text{ nS}$ | $E_{GIRK} = E_K$ | | | |
| $I_{Holo}$ | $g_{Holo} = 50 \text{ nS}$ | $\tau_{Holo} = 100 \text{ ms}$ | $E_{Holo} = E_{Syn}$ | | |

Activation (*m*_*i*_) and inactivation (*h*_*i*_) of voltage-dependent channels are described by the following differential equation:

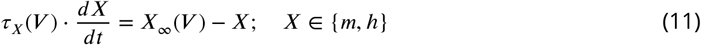

where *X*_oo_ represents steady-state activation/inactivation and *τ*_*X*_ is a time constant. For *I*_*Na*_, *I*_*NaP*_, and *I*_*Ca*_, the functions *X*_oo_ and *τ*_*X*_ take the forms

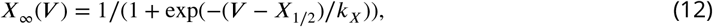

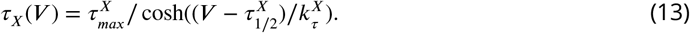

The values of the parameters (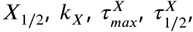 and 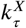)corresponding to *I*_*Na*_ *I*_*NaP*_ and *I*_*Ca*_ are given in Table 1.

For the voltage-gated potassium channel, the steady-state activation 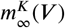 and time constant 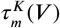 are given by the expressions

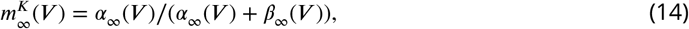

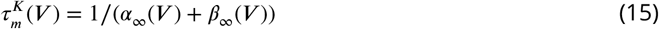

where

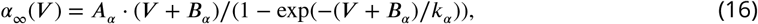

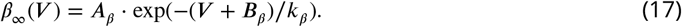

The values for the constants *A*_*α*_, *A*_*β*_, *B*_*α*_, *B*_*β*_, *k*_*α*_, and *k*_*β*_are also given in Table 1.

*I*_*CAN*_ activation depends on the *Ca*^2+^ concentration in the cytoplasm ([*Ca*]_*Cyto*_) and is given by:

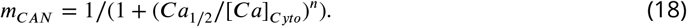

The parameters *Ca*_1/2_ and *n* represent the half-activation *Ca*^2+^ concentration and the Hill coefficient, respectively, and are included in Table 1.

The dynamics of [*Ca*]_*Cyto*_ is determined in part by the balance of *Ca*^2+^ effux toward a baseline concentration via the *Ca*^2+^ pump and *Ca*^2+^ influx through voltage-dependent activation of *I*_*Ca*_ and synaptically triggered *Ca*^2+^ transients, with a percentage (*P*_*SynCa*_) of the synaptic current (*I*_*Syn*_) carried by *Ca*^2+^ ions. Additionally, the model includes an intracellular compartment that represents the endoplasmic reticulum (ER), which impacts [*Ca*]_*Cyto*_. The ER removes *Ca*^2+^ from the cytoplasm via a sarcoplasmic/endoplasmic reticulum *Ca*^2+^-ATPase (SERCA) pump, which transports *Ca*^2+^ from the cytoplasm into the ER (*J*_*S RCA*_), and releases *Ca*^2+^ into the cytoplasm via calcium-dependent activation of the inositol triphosphae (IP3) receptor (*J*_*IP* 3_). Therefore, the dynamics of [*Ca*]_*Cyto*_ is described by the following differential equation:

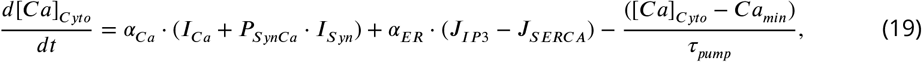

where *α*_*Ca*_ = 2.5 · 10^−5^ *mM* /*f C* is a conversion factor relating current to rate of change of [*Ca*]_*Cyto*_, *τ*_*pump*_ = 500 *ms* is the time constant for the *Ca*^2+^ pump, *Ca*_*min*_ = 5.0 · 10^−6^ *mM* is a minimal baseline calcium concentration, and *a*_*R*_ = 2.5 · 10^−5^ is the ratio of free to bound *Ca*^2+^ in the ER.

The flux of *Ca*^2+^ from the ER to the cytoplasm through the IP3 receptor is modeled as:

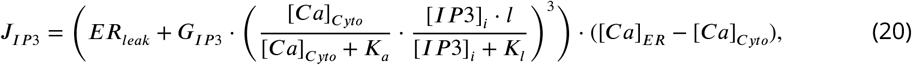

where *ER*_*leak*_ = 0.1/*ms* represents the leak constant from the ER stores, *G*_*IP* 3_ = 77, 500/*ms* represents the permeability of the IP3 channel, *K*_*a*_ = 1.0 · 10^−4^ *mM* and *K*_*l*_ = 1.0 · 10^−3^ *mM* are dissociation constants, and [*IP* 3]_*i*_ = 1.5 · 10^−3^ *mM* is the cytoplasm IP3 concentration. Finally, the *Ca*^2+^-dependent IP3 gating variable, *l*, and the *Ca*^2+^ concentration in the ER, [*Ca*]_*R*_, are determined by the following equations:

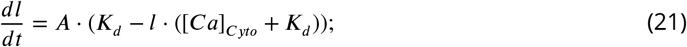

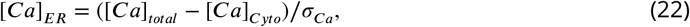

where *A* = 0.1 *mM* /*ms* is a conversion factor, *K*_*d*_ = 0.2 · 10^−3^ *mM* is the dissociation constant for IP3 inactivation, [*Ca*]_*total*_ is the total intracellular calcium concentration and *<Y*_*Ca*_ = 0.185 is the ratio of cytosolic to ER volume. The total intracellular calcium concentration is described as:

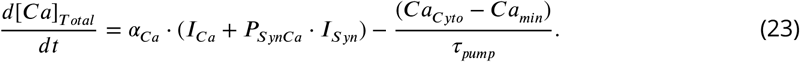

Finally, removal of *Ca*^2+^ from the cytoplasm by the SERCA pump is modeled as:

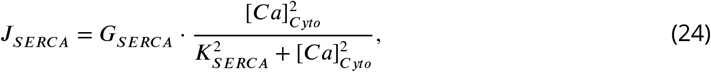

where *G*_*S RCA*_ = 0.45 *mM* /*ms* is the maximal flux through the SERCA pump, and *K*_*S RCA*_ = 7.5 · 10^−5^ *mM* is a dissociation constant.

When we include multiple neurons in the network, we can index them with subscripts. The total synaptic conductance (*g*_*Syn*_)_*i*_ of the *i*^*th*^ target neuron is described by the following equation:

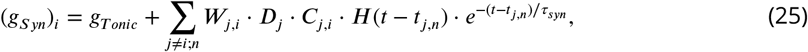

where *g*_*T onic*_ is a fixed or tonic excitatory synaptic conductance (e.g., from respiratory control areas outside of the preBötC) that we assume impinges on all neurons, *W*_*j,i*_ represents the weight of the synaptic connection from neuron *j* to neuron *i, D*_*j*_ is a scaling factor for short-term synaptic depression in the presynaptic neuron *j* (described in more detail below), *C*_*j,i*_ is an element of the connectivity matrix (*C*_*j,i*_ = 1 if neuron *j* makes a synapse with neuron *i* and *C*_*j,i*_ = 0 otherwise), *H*(.) is the Heaviside step function, and *t* denotes time. *τ*_*Syn*_ is an exponential synaptic decay constant, while *t*_*j,n*_ is the time at which the *n*^*th*^ action potential generated by neuron *j* reaches neuron *i*.

Synaptic depression in the *j*^*th*^ neuron (*D*_*j*_) was simulated using an established mean-field model of short-term synaptic dynamics (***Abbott et al., 1997***; ***Dayan and Abbott, 2001***; ***Morrison et al., 2008***) as follows:

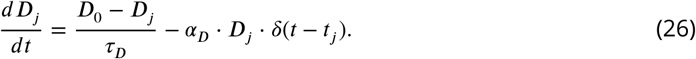

Where the parameter *D*_0_ = 1 sets the maximum value of *D*_*j*_, *τ*_*D*_ = 1000 *ms* sets the rate of recovery from synaptic depression, *α*_*D*_ = 0.2 sets the fractional depression of the synapse each time neuron *j* spikes and *δ*(.) is the Kronecker delta function which equals one at the time of each spike in neuron *j* and zero otherwise. Parameters were chosen to qualitatively match data from ***Kottick*** and Del Negro (***2015***).

When we consider a two-neuron network (Fig. 2), we take *W*_1,2_ = *W*_2,1_ = 0.006 and *C*_1,2_ = *C*_2,1_ = 1. For the full preBötC population model comprising rhythm and pattern generating subpopulations, the weights of excitatory conductances were uniformly distributed such that *W*_*j,i*_ = *U* (0, *W*_*Max*_) where *W*_*Max*_ is a constant associated with the source and target neurons’ populations; with each such pair, we also associated a connection probability and used this to randomly set the *C*_*j,i*_ values (see Table 3). Effects of opioids on synaptic transmission for source neurons in the rhythmogenic subpopulation (Fig. 6) were simulated by scaling *W*_*j,i*_ with the parameter *γ*_*µO R*_ which ranged between 0 and 0.5 and sets the percent synaptic block.

### Network construction

The relative proportions of neurons assigned to the rhythm and pattern generating preBötC sub-populations were chosen based on experimental data. For example, ***Kallurkar et al. (2020***) found that 20 ± 9% of preBötC inspiratory neurons are active during burstlets at *K*_*Bath*_ = 9 *mM*. Moreover, the rhythm and pattern generating neurons are hypothesized to be represented by the subsets of Dbx1 positive preBötC neurons that are somatostatin negative (*SST* ^−^) and positive (*SST* ^+^), respectively (***Cui et al., 2016***; ***Ashhad and Feldman, 2020***). Somatostatin positive neurons are estimated to comprise 72.6% of the *Dbx*1^+^ preBötC population (***Koizumi et al., 2016***). Therefore, our preBötC network was constructed such that the rhythm and pattern forming subpopulations represent 25% and 75% of the *N* = 400 neuron preBötC population (i.e., *N*_*R*_ = 100 and *N*_*P*_ = 300). The rhythm and pattern generating neurons are distinguished by their *I*_*NaP*_ conductances.

The synaptic connection probabilities within the rhythm (*P*_*RR*_ = 13%) and pattern (*P*_*P P*_ = 2%) generating neurons were taken from previous experimental findings (***Rekling et al. (2000***) and ***Ashhad and Feldman (2020***), respectively). The connection probabilities between the rhythm and pattern generating populations are not known and in the model were set at (*P*_*RP*_ = *P*_*P R*_ = 30%) such that the total connection probability in the network is approximately 13% (***Rekling et al., 2000***).

Heterogeneity was introduced by normally distributing the parameters *g*_*leak*_, *g*_*NaP*_ and *g*_*CAN*_ as well as uniformly distributing the weights (*W*_*j,i*_) of excitatory synaptic connections; see Tables 2 and 3. Additionally, *g*_*leak*_ was conditionally distributed with *g*_*NaP*_ in order to achieve a bivariate normal distribution between these two conductances, as suggested by ***Del Negro et al. (2002a***); ***Koizumi and Smith (2008***). In our simulations, this was achieved by first normally distributing *g*_*NaP*_ in each neuron according to the values presented in Table 2. Then used a property of bivariate normal distribution which says that the conditional distribution of *g*_*leak*_ given *g*_*NaP*_ is itself a normal distribution with mean 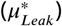 and standard deviation 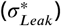 described as follows:

**Table 2.** Distributed channel conductances.

| Type | $g_{NaP} (nS)$ | | $g_{Leak} (nS)$ | | $g_{CAN} (nS)$ | |
| --- | --- | --- | --- | --- | --- | --- |
| | $\mu$ | $\sigma$ | $\mu$ | $\sigma$ | $\mu$ | $\sigma$ |
| Rhythm | 3.33 | 0.75 | $(K_{Bath} - 3.425)/4.05$ | $0.05 \cdot \mu_{leak}$ | 0.0 | 0.0 |
| Pattern | 1.5 | 0.25 | $(K_{Bath} - 3.425)/4.05$ | $0.025 \cdot \mu_{leak}$ | 2.0 | 1.0 |

**Table 3.**
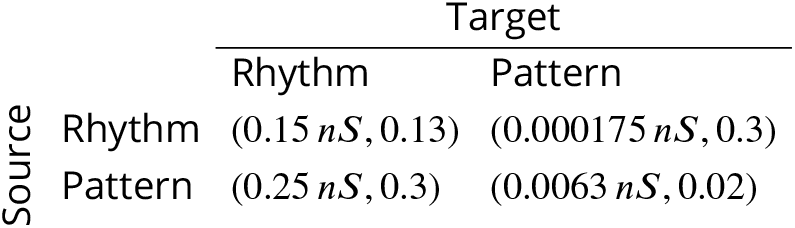
Maximal synaptic weights and connection probabilities between and within rhythm and pattern generating preBötC subpopulations (*W*_*Max*_, *P*).

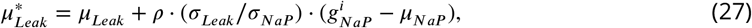

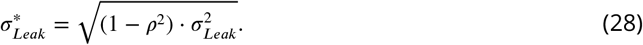

Where *µ*_*Leak*_ and *µ*_*NaP*_ are the mean and, *$σ*_*Leak*_ and *σ*_*NaP*_ are the standard deviation of the *g*_*Leak*_ and *g*_*NaP*_ distributions. Finally, *p* = represents the correlation coefficient and 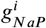 represents the persistent sodium current conductance for the *i*^*th*^ neuron. All parameters are given in Table 2.

### Activation dynamics of *I*_*Holo*_

Holographic stimulation was simulated by activating *I*_*Holo*_ in small sets of randomly selected neurons across the preBötC population. Activation of this current was simulated by the following equation:

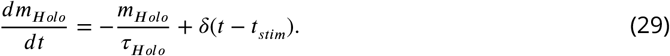

Where *m*_*Holo*_ represents the channel activation and ranges between 0 and 1, *τ*_*Holo*_ represents the decay time constant, and *δ* (.) is the Kronecker delta function which represents the instantaneous jump in *m*_*Holo*_ from 0 to 1 at the time of stimulation (*t*_*stim*_). Parameters were chosen such that the response in stimulated neurons matched those seen in ***Kam et al. (2013b***). All parameters are given in Table 1.

### Data analysis and definitions

Data generated from simulations was post-processed in Matlab (Mathworks, Inc.). An action potential was defined to have occurred in a neuron when its membrane potential *V*_*m*_ increased through −35*mV*. Histograms of population activity were calculated as the number of action potentials per 20 *ms* bin per neuron, with units of *AP s*/(*s* · *neuron*). Network burst and burstlet amplitudes and frequencies were calculated by identifying the peaks and the inverse of the interpeak interval from the population histograms. The thresholds used for burst and burstlet detection were 30 *spk*/*s*/*N* and 2.5 *spk*/*s*/*N*, respectively. For the simulated holographic stimulation simulations, the start of a network burst was defined as the time at which the integrated preBötC population activity increased through the threshold for burst detection, while the end of a network burst was defined as the time at which the integrated preBötC activity returned to exactly zero.

### Integration methods

All simulations were performed locally on an 8-core Linux-based operating system or on compute nodes at the University of Pittburgh’s Center for Research Computing. Simulation software was custom written in C++. Numerical integration was performed using the first-order Euler method with a fixed step-size (*t*) of 0.025*ms*. All model codes will be made freely available through the ModelDB sharing site hosted by Yale University upon publication of this work.

## Supplementary Material

**Figure 2-Figure Supplement 1.**
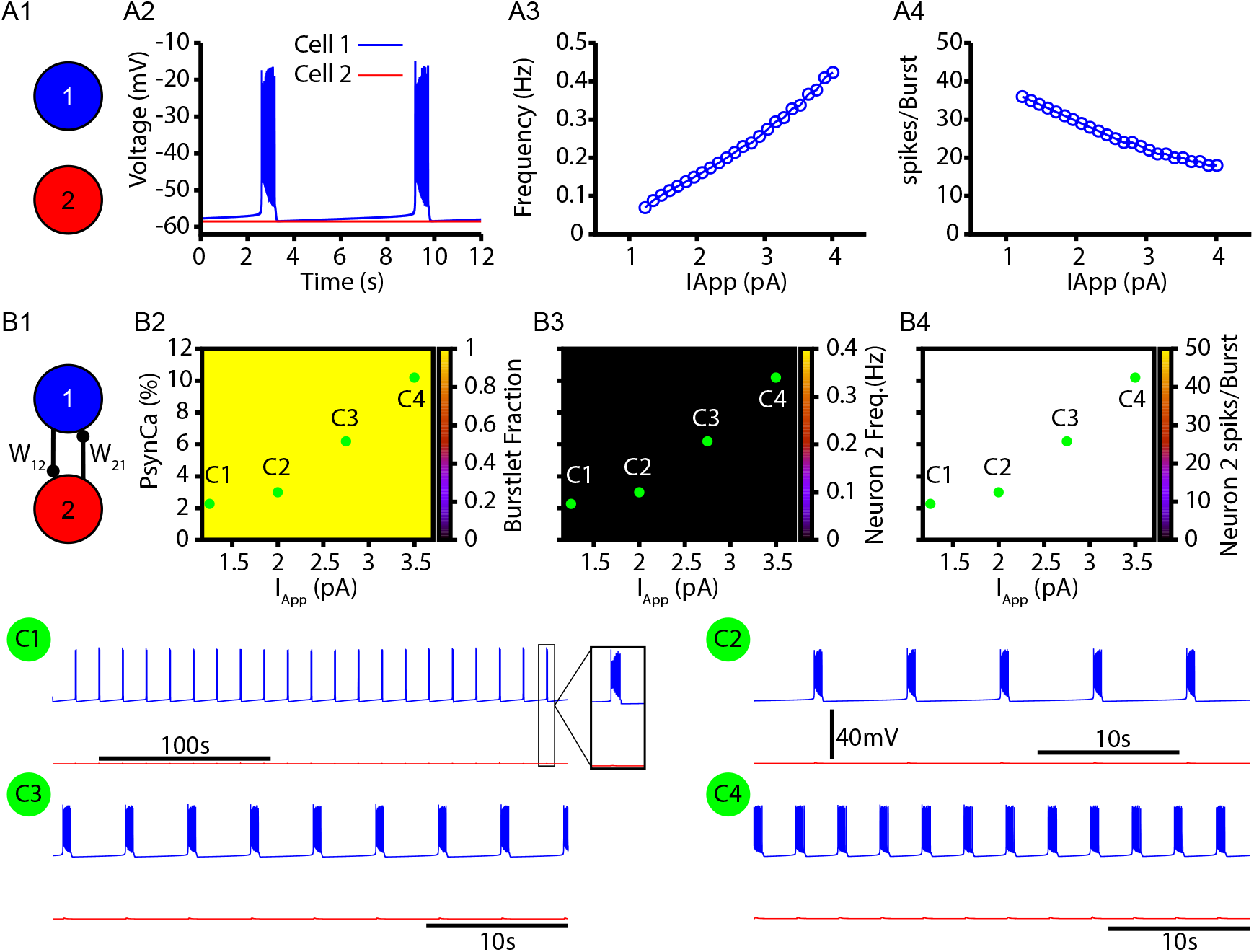
Without CICR, the two neuron network fails to generate bursts (recruitment of neuron 2). These simulations are identical to those in Fig. 2 except the conductance of the IP3 receptor is set to zero (*G*_*IP* 3_ = 0/*ms*). (A1) Schematic diagram of the synaptically uncoupled network. The rhythm and pattern generating components of the network are represented by neuron 1 and 2, respectively. (A2) Example trace showing intrinsic bursting in neuron 1 and quiescence in neuron 2. (A3) Burst frequency and (A4) the number of spikes per burst in neuron 1 as a function of an applied current (*I*_*AP P*_). Neuron 2 remained quiescent within this range of *I*_*AP P*_. (B1) Schematic diagram of the synaptically coupled network. (B2-B4) 2D plots characterizing the (B2) burstlet fraction, (B3) neuron 2 (burst) frequency, and (B4) neuron 2 spikes per burst (burst amplitude) as a function of *I*_*AP P*_ and *P*_*SynCa*_. (C1-C4) Example traces for neuron 1 and 2 for various *I*_*AP P*_ and *P*_*SynCa*_ values indicated in (B2-B4). Notice that neuron 2 is never recruited by the bursting in neuron 1 for any of the conditions tested. The model parameters used in these simulations are: (neuron 1 & 2) *K*_*Bath*_ = 8 *mM, g*_*Leak*_ = 3.35 *nS, W*_12_ = *W*_21_ = 0.006 *nS*; (Neuron 1) *g*_*NaP*_ = 3.33 *nS, g*_*CAN*_ = 0.0 *nS*, Neuron 2 *g*_*NaP*_ = 1.5 *nS, g*_*CAN*_ = 1.5 *nS*.

**Figure 3-Figure Supplement 1.**
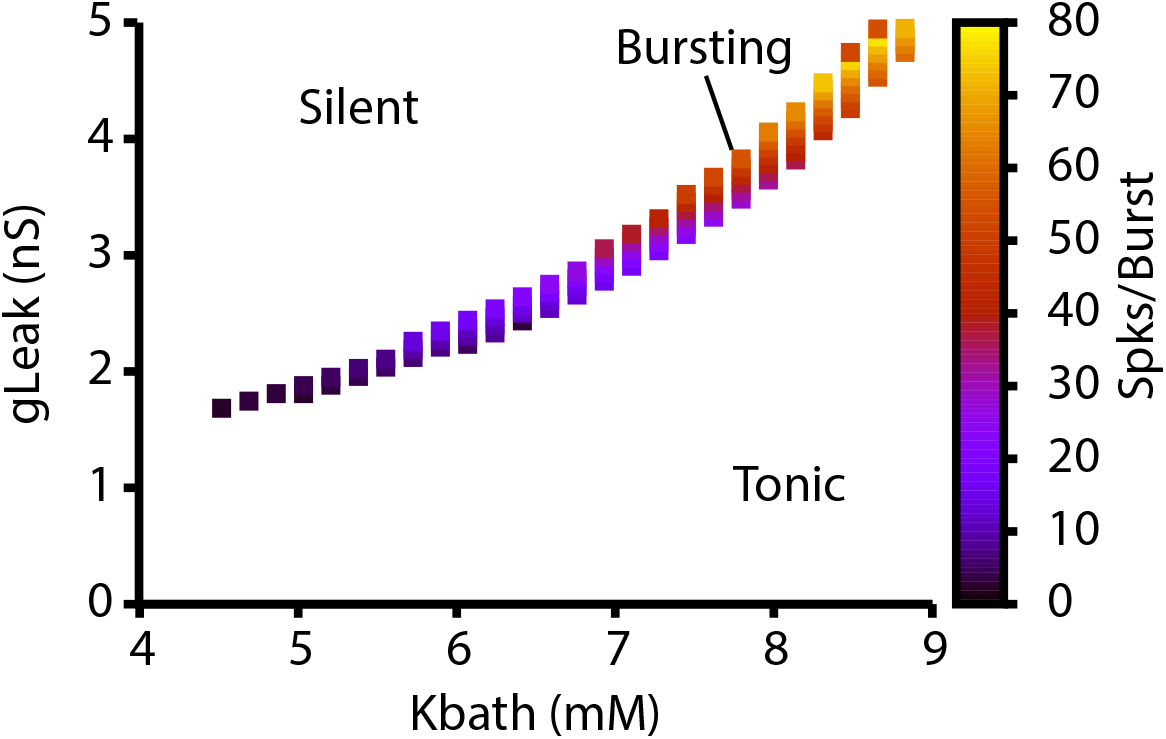
Dependence of intrinsic cellular dynamics and the number of spikes per burst on *K*_*bath*_ and *g*_*Leak*_. For these simulations *g*_*NaP*_ = 5.0 *nS*.

